# Pan-cancer transcriptional regulatory network analysis reveals key drivers and epigenetic modulators in tumorigenesis

**DOI:** 10.1101/2025.03.18.643959

**Authors:** Yaluan Yin, Junhua Zhang

## Abstract

Cancer remains a leading cause of mortality worldwide, largely due to the complexity of its transcriptional and epigenetic dysregulation. However, integrating multi-omics data to construct context-specific transcriptional regulatory networks (TRNs) and dissecting their pan-cancer relevance remain challenging. Here, we developed a framework integrating gene expression, chromatin accessibility, and DNA methylation data to reconstruct TRNs across 16 cancer types from The Cancer Genome Atlas. Using the PECA model, we identified TF-RE-TG (transcription factor-regulatory element - target gene) regulatory relationships and refined networks using protein-protein interaction or methylation-aware filtering. Our analysis revealed cancer-specific and pan-cancer dysregulated TFs and TGs. Notably, hub TFs like *TBP* and *SP2* orchestrated pan-cancer modules linked to DNA repair. Survival analysis identified prognostic TGs (e.g., *TAS2R14, ANLN*), while enhancer and methylation analyses highlighted epigenetic-related regulatory axes (e.g., *NFIB-RPS6KB2, MYB-BTK*) in tumorigenesis. Our study provides a comprehensive resource of cancer TRNs, unveiling conserved regulatory mechanisms and epigenetic vulnerabilities, with implications for precision oncology.

## 1 Introduction

Cancer remains one of the most formidable threats to human health, characterized by its complex pathogenesis and high mortality rates. Despite advancements in oncology, the lack of effective therapeutic strategies underscores the urgent need to unravel the molecular mechanisms driving tumorigenesis and progression. Systematic analysis of high-throughput multi-omics data may offer unprecedented insights into cancer biology and precision medicine. Large-scale initiatives such as The Cancer Genome Atlas (TCGA) [1], the International Cancer Genome Consortium (ICGC) [2], and the Cancer Cell Line Encyclopedia (CCLE) [3] have generated genomics, transcriptomics, and epigenomics data, laying the foundation for understanding cancer heterogeneity and identifying therapeutic targets.

Central to cancer biology is the dysregulation of transcriptional control mechanisms. Transcriptional regulation, governed by transcription factors (TFs) and epigenetic modifications, dictates spatiotemporal gene expression patterns critical for maintaining cellular homeostasis. Aberrant TF activity usually induces abnormal biological processes, such as uncontrolled proliferation, immune evasion, and drug resistance, which are hallmarks of cancer [4–6], highlighting TF’s pivotal role in disease pathogenesis. Concurrently, epigenetic dysregulation, including DNA methylation and chromatin remodeling, further disrupts transcriptional programs. For instance, promoter hypermethylation silences tumor suppressor genes, while gene body methylation may activate oncogenes [7]. Chromatin accessibility, a dynamic epigenetic feature, governs TF binding and transcriptional activity, with alterations observed in glioblastoma, lung cancer and ovarian cancer [8–10]. The ATAC-seq data from TCGA samples has mapped chromatin accessibility landscapes across 23 cancer types [11], underscoring its relevance to tumor biology.

Reconstructing transcriptional regulatory networks (TRNs) is essential for decoding the interplay between TFs, regulatory elements (REs), and target genes (TGs). Reverse engineering approaches, such as correlation-based models [12], regression models [13], probabilistic frameworks [14], Bayesian network-based methods [15], and Transformer-based deep learning models [16, 17], leverage gene expression data to infer TF-TG interactions. Experimental methods like ChIP-seq [18] may provide direct evidence of TF binding, yet their high cost and cell-type specificity limit scalability. Multi-omics integration has emerged as a powerful strategy, combining transcriptomic and epigenomic data to enhance TRN accuracy. For example, PECA [19], SCENIC+ [20] and LINGER [21] utilize chromatin accessibility and gene expression to identify cis-regulatory elements [22], while methylation-driven TRNs [23] reveal cancer-specific epigenetic dependencies. Despite progress [24, 25], existing methods often neglect the synergistic effects of REs and DNA methylation, limiting their ability to capture the full complexity of cancer regulatory landscapes. Furthermore, enhancers are distal cis-regulatory DNA elements, typically located in open chromatin regions, that can activate or enhance gene transcription. The regulatory role of enhancers is closely related to the occurrence and development of cancer, with most cancers being controlled by enhancers or super-enhancers [26]. Comprehensively investigating the interplay between REs, enhancer elements, and DNA methylation in transcriptional dysregulation for pan-cancer will be of great significance for deepening the understanding of cancer pathogenesis and progression, but such research is very scarce.

In this study, we addressed these gaps by constructing pan-cancer TRNs through integrative analysis of gene expression (RNA-seq), chromatin accessibility (ATAC-seq), and DNA methylation data from TCGA. Our approach extended the PECA model [19] to incorporate protein-protein interaction (PPI) networks or methylation profiles to refine regulatory relationships. Applied to 16 TCGA cancer types, we investigated the biological processes that the differentially expressed TFs and TGs in the constructed TRNs were involved in, and elucidated their functional roles in tumorigenesis. We validated candidate prognostic biomarkers (e.g., *TAS2R14, ANLN*) through survival analysis. Notably, we identified conserved regulatory modules (e.g., *TBP/SP2* hubs) and cancer-specific subnetworks driving oncogenic processes. Furthermore, we explored the interplay between REs, enhancer elements, and CpG methylation, providing mechanistic insights into enhancer-mediated and methylation-sensitive transcriptional dysregulation. By benchmarking against established resources like BART Cancer [27] and Cistrome Cancer [28], we highlighted conserved (e.g., the *FOXM1/E2F1* -hub module in bladder urothelial carcinoma) and novel regulatory modules with therapeutic potential.

This work advances cancer research by unifying multi-omics data to reconstruct context-specific TRNs, thus bridging the gap between epigenetic modifications and transcriptional dysregulation. Our findings not only illuminate key drivers of tumor progression but also offer a framework for precision oncology, where targeting critical regulators or genes could disrupt oncogenic networks and improve patient outcomes.

## 2 Materials and methods

### 2.1 Data sources and preprocessing

In this study pan-cancer transcriptional regulatory networks were constructed using multi-omics data from TCGA, including RNA-seq, ATAC-seq, DNA methylation data, et al. The used cancer types and their abbreviations are listed in Table S1 (Supplementary Material 1). RNA-seq data was download from TCGA database (https://portal.gdc.cancer.gov/), and raw counts for protein-coding genes (GENCODE annotation [29]) were filtered to retain genes expressed in 50% of samples across 16 cancers. Count data were log-transformed for downstream analysis. ATAC-seq data from the chromatin accessibility atlas [11] were normalized per sample. DNA methylation *β*-values were obtained from the TCGA Infinium HumanMethylation450k platform, focusing on hypomethylated regulatory elements (REs; *β <* 0.3) to prioritize TF binding events. TF motif information was integrated from the NCBI [30], UCSC Genome Browser [31], and JASPAR [32] databases. The PPI network was from STRING [33].

### 2.2 Construction of transcriptional regulatory networks

The PECA model [19] was employed to infer TF-RE-TG regulatory relationships. This model simulates target gene expression through three probabilistic equations: (1) recruitment of chromatin regulators (CRs) to REs based on TF-CR interactions (Eq. 1), (2) RE activation influenced by recruited CRs (Eq. 2), and (3) TG transcription driven by active RE-bound TFs (Eq. 3).

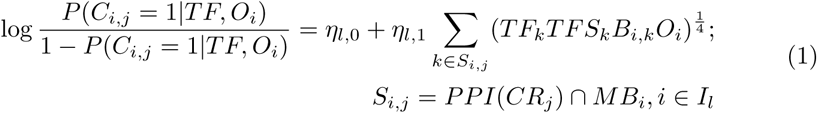

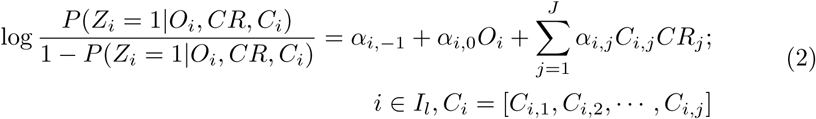

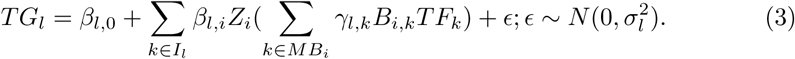

Here, *TF_k_*, *CR_j_* and *TG_l_* represent the expression values of the *k*-th TF, the *j*-th CR and *l*-th TG, respectively; *TFS_k_* denotes the tissue specificity score of the *k*-th TF’s expression; *O_i_* indicates the degree of chromatin accessibility of the *i*-th RE; *Z_i_* and *C_i,j_* are the indicative functions of whether the *i*-th RE is active and whether the *j*-th CR is recruited to the *i*-th RE, respectively; *PPI*(*CR_j_*) represents the subset of TFs known to interact with the *j*-th CR; *MB_i_* represents the TF subset with significant motif matches in the *i*-th RE; while *B_i,k_* represents the sum of *−log*(*p*-values) of the *k*-th TF’s motifs in the *i*-th RE.

For each target gene, the PECA model (Eqs. 1 - 3) selects REs whose corresponding coefficient *β* is nonzero and TFs whose coefficient *γ* is nonzero, and outputs the regulatory relations RE-TGs, TF-TGs, and TF-RE-TGs. Preliminary networks were intersected with the STRING PPI network [33] to enhance reliability.

Hypomethylated motifs were identified using HOMER [34], and corresponding REs were retained to refine TF-RE-TG networks.

### 2.3 Differential expression and functional enrichment

Differentially expressed TFs and TGs were identified by comparing cancer and normal samples (FDR *<* 0.05). Functional enrichment analysis was performed via DAVID [35] for Gene Ontology (GO) terms and KEGG pathways.

### 2.4 Integration with external databases

Overlaps with BART Cancer [27] and Cistrome Cancer [28] were to identify high-confidence regulatory modules. For enhancer analysis, cancer-related enhancers from the HEDD database [36] were mapped to REs.

### 2.5 Prognostic modeling and survival analysis

TGs with low methylation modification in the promoter region are often highly expressed. These TGs are of particular interest due to their potential association with cancer progression. Here we pay more attention to such TGs for prognostic analysis. Univariate Cox regression (Eq. 4) was applied to significantly upregulated TGs to identify prognosis-related candidates:

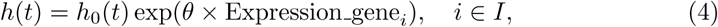

where *h*(*t*) represents survival time, and *I* denotes the set of significantly overexpressed TGs. Genes with non-zero coefficient *θ* were selected as candidate TGs. Usually, hazard ratio (HR) is defined as *e^θ^*.

Further, to reduce false positives, we conducted LASSO-COX regression (Eq. 5) on the candidate TGs:

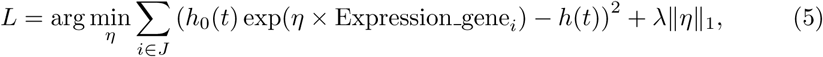

where *J* represents the set of candidate TGs, and *λ* is the regularization parameter. Now the genes with non-zero coefficient *η* were considered as associated TGs with the prognosis of the cancer.

Risk scores (Eq. 6) were calculated for risk assessment of cancer patients:

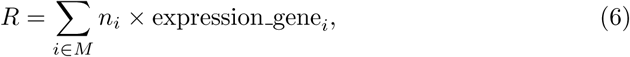

where *M* represents prognosis-associated genes and *η_i_* denotes regression coefficients obtained in Eqs. 4 and 5. Patients were stratified into high- and low-risk groups, and survival differences were assessed via Kaplan-Meier analysis [37].

### 2.6 DNA methylation and enhancer analysis

An enhancer is a distal cis-regulated DNA element, usually located in the region of chromatin accessibility, that can activate or enhance transcription of genes. The regulation of enhancer closely affects the initiation and progression of cancer. Most oncogenes are controlled by enhancer or super enhancer [26]. In this study enhancers overlapping with REs were analyzed for TF-TG consistency with HEDD predictions. Some important multiple regulation relationships TF-RE(-enhancer)-TGs were identified.

DNA methylation can affect the structure of chromatin and participate in gene expression regulation. Generally, DNA methylation in gene promoter regions is thought to inhibit the transcription process. Thus when constructing regulatory networks by further incorporating DNA methylation data, we focus on TF’s motifs with hypomethylated CpG sites (*β <* 0.3), and then link the REs to TF banding and TG dysregulation. Based on this method, methylation-driven regulatory modules (TF-RE(-mCpG)-TGs) (where mCpG represents the methylated CpG site on DNA) were prioritized for functional studies.

This integrative approach enables systematic exploration of pan-cancer transcriptional regulation, combining multi-omics data with robust statistical models to uncover drivers of cancer progression and prognosis.

## 3 Results

First, by incorporating PPI networks with the PECA model, we reconstructed pan-cancer TRNs based on RNA-seq and ATAC-seq for 16 cancer types from TCGA (including BLCA, BRCA, COAD, ESCA, HNSC, KIRC, KIRP, LGG, LIHC, LUAD, LUSC, PRAD, SKCM, STAD, THCA, and UCEC) (Supplementary Material 1, Table S1). Then we further extend the PECA model to incorporate methylation profiles to uncover more refined regulatory relationships (now for 15 cancer types because UCEC has no DNA methylation data for samples with both RNA-seq and ATAC-seq).

### 3.1 Cancer transcriptional regulatory networks based on RNA-seq, ATAC-seq and the PPI network

In this section we constructed pan-cancer transcriptional regulatory atlas for 16 cancer types from TCGA. Based on differential expression analysis of cancer vs normal samples, the key questions are: 1) What functions do the significantly over/underexpressed TFs possess? And what downstream genes do they regulate? 2) What biological processes are the significantly differentially expressed TGs primarily involved in? And by what functional TFs are they regulated? Trying to answer these questions, we explored TFs and TGs of significant importance from both cancer-specific and pan-cancer perspectives, and studied important regulatory relationships in cancer by analyzing the regulatory subnetworks or modules they participate in.

#### 3.1.1 Dysregulated TFs drive shared and cancer-specific regulatory mechanisms

Here we take some typical similar cancers as examples to focus on their comparative analysis. The analysis of gynecological cancers (BRCA and UCEC) and lung cancers (LUAD and LUSC) revealed commonalities and specificities in dysregulated TFs and their downstream pathways.

##### Significantly underexpressed TFs and their regulated subnetworks

**BRCA and UCEC:** These are two typical gynecological tumors, and previous studies have shown that they share some similarities. Here, we compare and analyze these two cancers from the perspective of transcriptional regulatory networks. We found that many significantly underexpressed TFs in BRCA and UCEC are the same, with more than half of the TFs in UCEC also appearing in BRCA, indicating some commonalities between UCEC and BRCA. Further, based on DAVID enrichment analysis [35], we examined the functions significantly enriched by underexpressed TFs in their respective cancers (Fig. 1). It can be seen that many typical cancer-related biological processes are common for these two types of tumors, such as angiogenesis, regulation of cell proliferation, cellular response to hypoxia, and negative regulation of cell apoptotic process. Additionally, BRCA also enriches processes like cellular response to growth hormone stimulation and the Notch signaling pathway, while UCEC significantly enriches processes like cellular response to estrogen and estradiol. These findings suggest that the significantly underexpressed TFs in BRCA and UCEC are closely related to the occurrence and development of cancer.

**Fig. 1:**
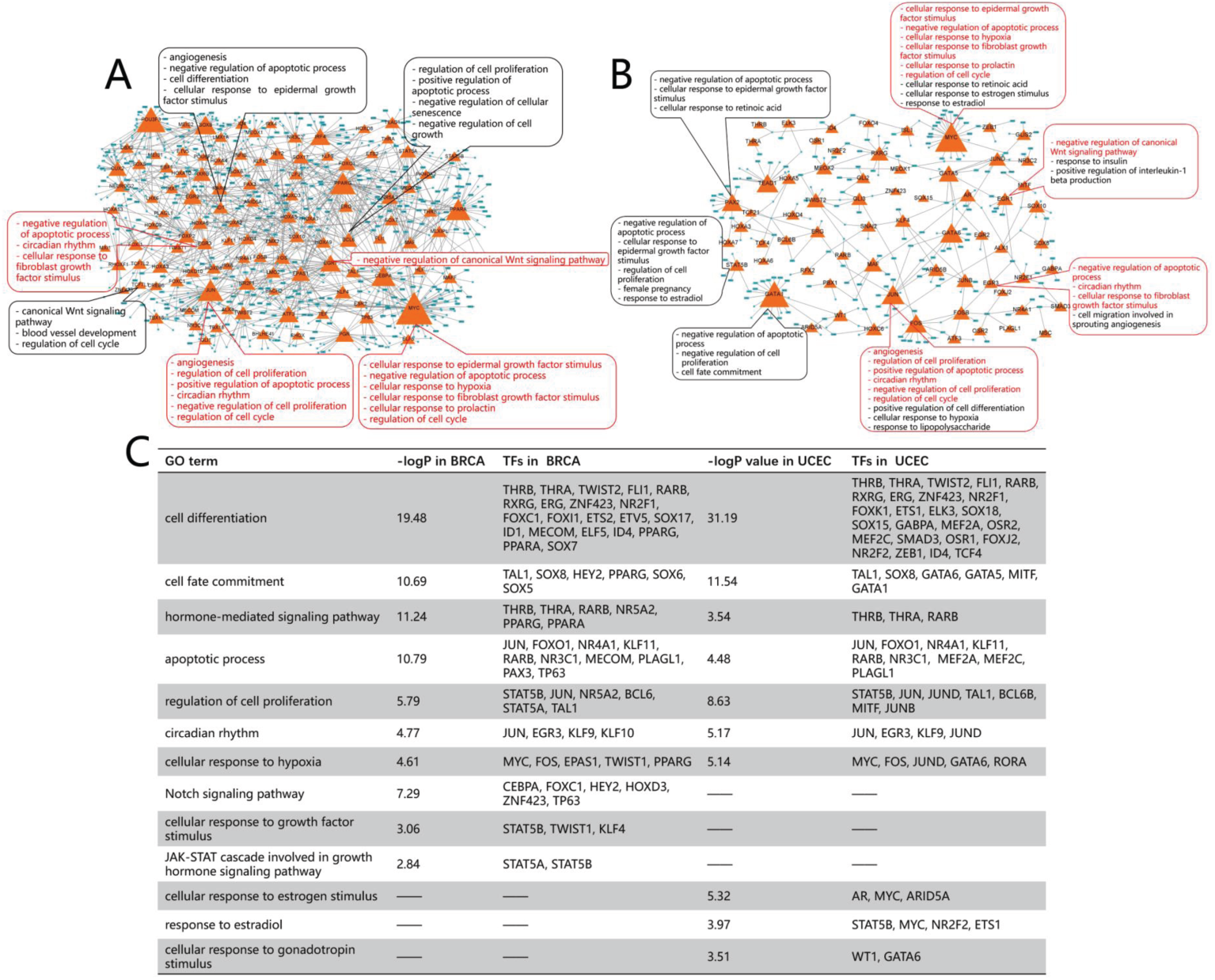
The enriched functions and regulatory subnetworks of significantly underexpressed TFs in BRCA and UCEC. The orange triangle and cyan rectangle represent TF and TG, respectively. The size of TF represents the number of regulatory relations. The text box shows the significantly enriched GO terms of the TF. The boxes and text highlighted in red represent the common enriched functions of TFs in both cancers. A: BRCA; B: UCEC; C: Comparison of significantly enriched functions of TFs in BRCA and UCEC.

Further, we examined the subnetworks regulated by significantly underexpressed TFs in both cancer types. We found that the TGs in the BRCA regulatory subnetwork are more numerous than those in UCEC. Enrichment analysis of TGs showed that BRCA TGs significantly enrich more KEGG signaling pathways. However, many of these pathways are the same in both cancers, such as the PI3K-Akt signaling pathway, MAPK signaling pathway, apoptosis, and Ras signaling pathway. This led us to wonder: Are the TFs regulating the same KEGG pathways the same in both cancers? And what are the similarities and differences in the biological processes they participate in? Here, we take two typical signaling pathways as examples: the RAS signaling pathway and the MAPK signaling pathway, and display the connected subnetworks of the regulatory relationships corresponding to these pathways (Fig. 2A-D).

The RAS oncogene is a typical oncogene in human tumors, and RAS gene mutations are common in both hematologic and solid tumors. Targeting the Ras signaling pathway is an important approach to overcoming tumors. Although the TFs regulating the Ras signaling pathway in BRCA and UCEC are almost completely different (Fig. 2A, B), they are involved in many of the same biological processes, such as negative regulation of cell proliferation, positive regulation of apoptotic process, cell differentiation, and angiogenesis. The downregulation of TFs weakens these biological functions, leading to abnormal proliferation and spread of cancer cells that should have undergone apoptosis or senescence, ultimately inducing cancer. Similar phenomena are observed in the MAPK signaling pathway (Fig. 2C, D). The MAPK signaling pathway transmits extracellular signals to the intracellular environment through a three-tier kinase cascade (MAPKKK *→* MAPKK *→* MAPK), regulating cell proliferation, differentiation, apoptosis, inflammatory responses, and vascular development. It is one of the most classic cancer signaling pathways. By comparing Fig. 2C and D, we see that TFs in both cancers participate in some common cancer-related biological processes, such as cell fate commitment, angiogenesis, cell differentiation, and cell proliferation.

**LUAD and LUSC:** For these two typical non-small-cell lung cancers, most of the significantly underexpressed TFs in LUAD were also identified in LUSC. These TFs significantly enriched many of the same biological functions in both cancers, such as angiogenesis, circadian rhythm, and the Notch signaling pathway, all of which are closely related to cancer development.

On the other hand, the TGs regulated by these differentially underexpressed TFs in LUAD and LUSC also significantly enriched many of the same KEGG pathways, such as the NF-*κ*B pathway, transcriptional misregulation in cancer, and the MAPK signaling pathway.

The NF-*κ*B pathway is closely related to cancer. Its persistent activation can lead to abnormal activity of cell cycle-related proteins, increase levels of cytokines that promote tumor growth, promote tumor tissue infiltration and metastasis, and enhance the anti-apoptotic ability of cancer cells. Among the TFs regulating the NF-*κ*B pathway in LUAD and LUSC, they share participation in cell differentiation, negative regulation of cell proliferation, and cellular response to hypoxia, all of which play crucial roles in cancer proliferation, invasion, and metastasis, and are key factors in cancer progression (Supplementary Material 1, Fig. S1A, B). Additionally, TFs in LUAD participate in the positive regulation of apoptosis, while TFs in LUSC participate in signal transduction and cellular response to transforming growth factor *β* stimulation (TGF-*β* signaling), all of which are closely related to cancer development.

For transcriptional misregulation, the TFs regulating this pathway in LUAD and LUSC participate in common cancer-related biological processes, including angiogenesis, cell differentiation, cellular response to hypoxia, and circadian rhythm, all of which significantly impact cancer development (Supplementary Material 1, Fig. S1C, D). Furthermore, TFs in LUAD also participate in cellular senescence and positive regulation of apoptosis, while TFs in LUSC participate in signal transduction. In this context, the downregulation of TFs leads to the loss of their corresponding functions, allowing cancer cells to proliferate indefinitely, further exacerbating cancer development.

**Fig. 2:**
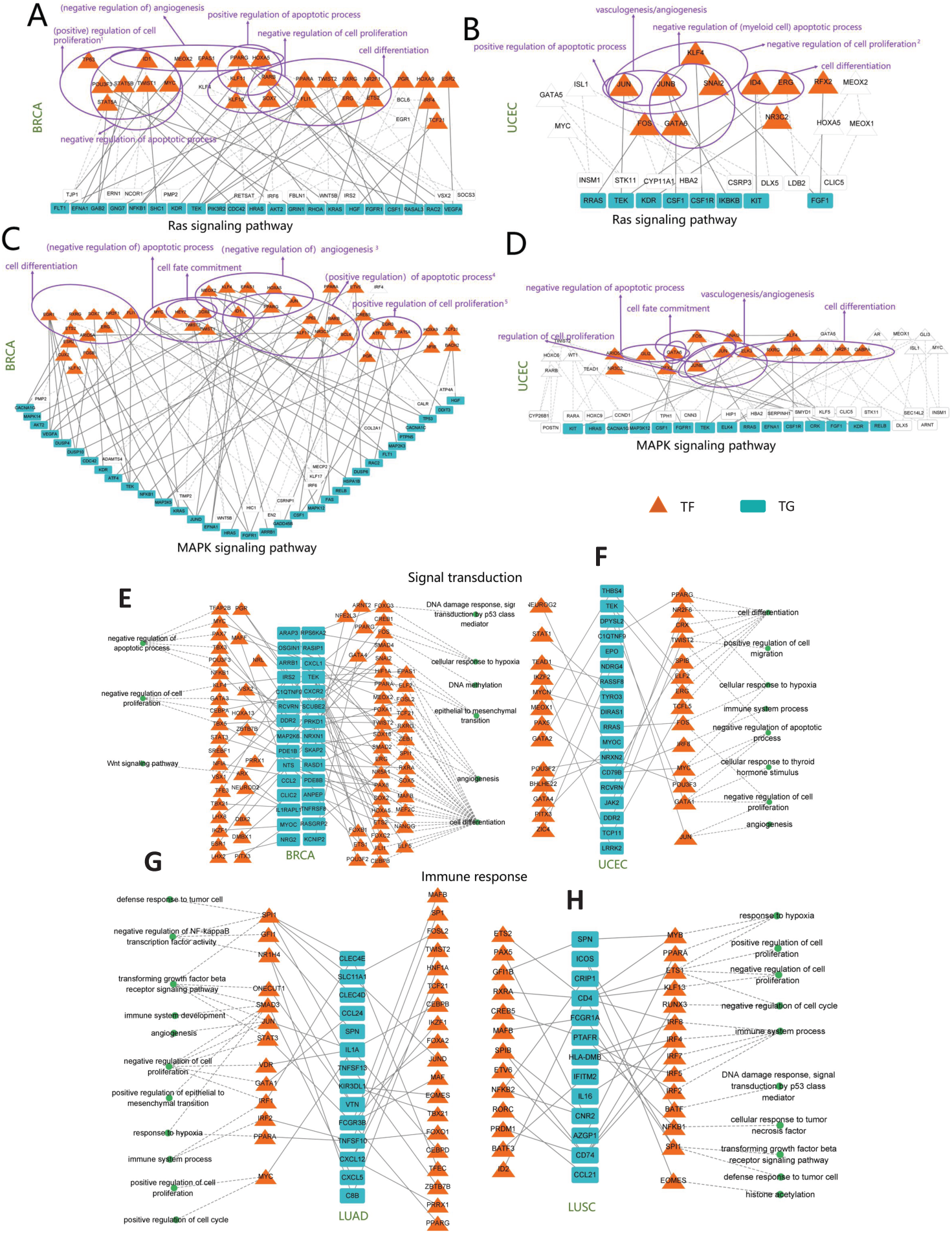
A-D: The functions and regulatory subnetworks of underexpressed TFs. A, B: TGs belong to Ras signaling pathway (A: BRCA, B: UCEC); C, D: TGs belong to MAPK signaling pathway (C: BRCA, D: UCEC). E-H: Functional subnetworks of TFs regulating underexpressed TGs in cancers. E, F: TGs are involved in signal transduction (E: BRCA, F: UCEC); G, H: TGs are related to immune response (G: LUAD, H: LUSC). The purple texts in A-D and the green dots in E-H mean GO terms. The white triangles and rectangles are other TFs and TGs added to make the subnet connected in A-D. To save space, we have placed the ‘Note’ about A-D into the ‘Supplementary Material 1’.

In summary, based on the results from BRCA and UCEC, as well as LUAD and LUSC, we find that the subnetworks regulated by significantly underexpressed TFs share many common or similar regulatory roles in similar cancers, but also have some differences, all of which are closely related to cancer development and worthy of further in-depth research and analysis.

##### Significantly overexpressed TFs and their regulated subnetworks

We also compared the subnetworks regulated by significantly overexpressed TFs in BRCA and UCEC, as well as LUAD and LUSC. Similar to the subnetworks regulated by underexpressed TFs, there are many common significantly overexpressed TFs between BRCA and UCEC, as well as between LUAD and LUSC. These TFs participate in similar functions in both cancers, significantly enriching many of the same biological functions that play roles in cancer (Supplementary Material 1, Fig. S2), such as cell differentiation, cell fate commitment, positive regulation of cell proliferation, negative regulation of apoptosis, glucose homeostasis, and immune system processes. However, compared to the underexpressed TFs, we still identified some specific biological processes involved by significantly overexpressed TFs (Supplementary Material 1, Fig. S2). For example, overexpressed TFs participate in epithelial-mesenchymal transition (EMT) (in LUSC) and mesenchymal-epithelial transition (MET) (in BRCA). This suggests that in LUSC, the overexpression of TFs such as *TRIM2B, LEF1, SNAI2*, and *SOX9* may cause epithelial cells to lose their epithelial phenotype and acquire a mesenchymal phenotype with high migration, invasion, antiapoptotic, and extracellular matrix degradation capabilities, leading to the migration and invasion of malignant cells derived from epithelial cells, accelerating cancer progression. In BRCA, the overexpression of TFs such as *WT1, GATA3*, and *PAX2* may reverse the EMT transformation of cancer cells, restoring the epithelial phenotype and adhesion ability, facilitating the ‘homing’ and proliferation of cancer cells, and forming metastatic foci.

Now we are further curious: How do significantly overexpressed TFs in cancer regulate downstream pathways? We explored this question in BRCA and UCEC using two typical KEGG signaling pathways (PI3K-Akt signaling pathway and MAPK signaling pathway) as examples.

The dysregulation of the PI3K-Akt signaling pathway is common in cancer. This pathway promotes cell metabolism, proliferation, survival, and growth. Analysis shows that significantly overexpressed TFs regulating this signaling in both BRCA and UCEC participate in biological processes such as cell proliferation and cell differentiation (Supplementary Material 1, Fig. S3A, C). Additionally, they each have unique characteristics. For example, in BRCA, the TF *E2F1* participates in the p53-mediated intrinsic apoptotic signaling pathway, while in UCEC, some TFs participate in the TGF-*β* signaling pathway.

For the MAPK signaling pathway, the overexpressed TFs regulating it in both BRCA and UCEC participate in important biological processes such as negative regulation of apoptosis, cell fate commitment, cell differentiation, and positive regulation of cell proliferation (Supplementary Material 1, Fig. S3B, D). In addition, the TFs in these two cancers also participate in some unique biological processes. Notably, MAPK signaling is the common enriched pathway of the TGs in BRCA and UCEC whether TFs overexpressed or underexpressed (Fig. 2C, D; Fig. S3B, D). Although the TFs are different in these two cases, they participate in important biological processes such as negative regulation of apoptosis, cell fate commitment, cell differentiation, and positive regulation of cell proliferation, all of which influence cancer migration and accelerate cancer progression.

#### 3.1.2 Dysregulated TGs highlight core cancer pathways and immune dysfunction

In the previous subsection we examined the significantly differentially expressed TFs and their regulated subnetworks in the constructed TRNs. Here we explore the biological processes that significantly differentially expressed TGs are involved in, and investigate the functions of the TFs regulating them. We will still take BRCA vs. UCEC and LUAD vs. LUSC as examples.

##### Significantly underexpressed TGs and regulatory TF subnetworks

We identified significantly underexpressed TGs in BRCA (247 genes), UCEC (138 genes), LUAD (170 genes), and LUSC (254 genes). Only 18 common TGs were shared between BRCA/UCEC and between LUAD/LUSC. Despite this small overlap, TGs were significantly enriched in common cancer-related biological functions. In BRCA/UCEC, enriched functions included signal transduction and positive regulation of inflammatory response. In LUAD/LUSC, enriched functions included immune response, positive regulation of phagocytosis, and endocytosis. Downregulation of these TGs disrupts cellular mechanisms, impairing processes like phagocytosis of cancer cells, ultimately affecting cancer development. Significantly enriched functions across all four cancers also included cancer-related processes like positive regulation of tumor necrosis factor, negative regulation of protein phosphorylation, and T cell differentiation. Weakening of these functions prevents cancer cell elimination, promoting tumorigenesis.

Examining TFs regulating these underexpressed TGs revealed key insights. For ‘signal transduction’ (BRCA/UCEC), the regulating TFs participated in shared processes including cell differentiation, cellular response to hypoxia, positive regulation of cell proliferation, negative regulation of apoptosis, and cellular responses to tumor necrosis factor, estrogen, estradiol, and progesterone (Fig. 2E, F). These functions collectively promote tumor growth, metastasis, and poor prognosis. For ‘immune response’ (LUAD/LUSC), the regulating TFs participated in immune-related processes like immune system processes, tumor defense response, and negative regulation of inflammatory response (Fig. 2G, H). Their involvement enhances cancer metastasis and poor prognosis.

##### Significantly overexpressed TGs and regulatory TF subnetworks

Significantly overexpressed TGs showed slightly larger overlaps: 23 common TGs between BRCA/UCEC and 28 between LUAD/LUSC. Overexpressed TGs in all four cancers were consistently enriched in cancer-related functions. BRCA/UCEC TGs were involved in DNA repair, apoptosis, cell proliferation, and cell cycle. LUAD/LUSC TGs were also enriched in DNA repair C a critical response to DNA damage. Impaired DNA repair can lead to cancer. Analysis of TFs regulating ‘DNA repair’ in all four cancers revealed similarities (e.g., *E2F, MYB, FOX, MAF* families, *TBP, TRIM28, HSF1*). These TFs participated in shared biological processes like cell differentiation, positive regulation of cell proliferation, cellular response to hypoxia, DNA damage response, and p53-mediated signal transduction (Supplementary Material 1, Fig. S4), all closely linked to cancer.

##### Further analysis in BRCA

Comparing biological processes enriched by underexpressed and overexpressed TGs in BRCA revealed only one common process: the MAPK signaling pathway. Over-expressed TGs showed specific enrichments; for example, 4 genes were involved in homologous recombination, a key DNA damage repair mechanism. Failure of this mechanism can severely damage cells and lead to cancer. Underexpressed TGs included 8 genes involved in the PPAR signaling pathway (regulating cell growth, differentiation, and apoptosis via lipid metabolism sensors) and 8 genes involved in the regulation of adipocyte lipolysis. Cancer often relies heavily on fat decomposition for growth.

TFs regulating both underexpressed and overexpressed TGs participated in many common processes (e.g., cell differentiation, regulation of cell proliferation, response to estrogen). Key differences were noted: TFs for underexpressed TGs participated in negative regulation of angiogenesis and cell growth, while TFs for overexpressed TGs participated in positive regulation of cell migration and DNA methylation. Crucially, the processes involving TFs regulating either underexpressed or overexpressed TGs play vital roles in cancer development and warrant further investigation.

#### 3.1.3 Pan-cancer validation and novel regulatory relationships

By benchmarking against established resources like BART Cancer (providing over/underexpressed gene sets and regulating TFs, but not specific TF-TG pairs) [27] and Cistrome Cancer (predicting targets for 575 TFs) [28], we highlight conserved and novel regulatory modules with therapeutic potential.

##### Pan-cancer validation by BART Cancer and Cistrome Cancer

We first took the intersection of differentially expressed TGs and corresponding TFs in our results with differentially expressed TGs and significantly ranked (*p*-value *<* 0.05) TFs in BART Cancer (Fig. 3A, B). Then we mapped TF-TG relationships from our network onto these intersections (Fig. 3C, D). We focused on cancers with abundant relationships, building subnetworks. Finally, we intersected these subnetworks with Cistrome Cancer to get regulatory modules or conserved relationships supported by multiple studies.

From the modules we found some key oncogenic drivers. For example, in BRCA, Module 1 (*ATF3, EGR1, CSRNP1, SOCS3*) promoted proliferation and apoptosis resistance (Fig. 4A). In fact, *ATF3* is an important TF and its overexpression enhances radiation/endocrine resistance in breast cancer [38]. *ATF3*’s targets (*CSRNP1, SOCS3*) inhibit apoptosis, synergizing with *ATF3* to promote proliferation. Further, *ATF3-EGR1* interaction contextually influences inflammation and tumor suppression/promotion [39]. These mean Module 1 likely induces breast cancer. While for Module 2 (*FLI1, GIMAP6, GIMAP8, S1PR1, TEK, VWF*) (Fig. 4A), previous studies have shown that *FLI1* can drive tumor metastasis and is an effective predictor of poor prognosis in breast cancer [40]. More than that, it has been found that in breast cancer, many important immune-related features are positively correlated with the expression level of *FLI1* [41], and its two TGs (*GIMAP6* and *GIMAP8*) are also members of the immune-related protein (GIMAP) family. It indicates that FLI1 may jointly participate in the immune system processes of cancer cells by regulating these two TGs. On the other hand, the other two TGs of *FLI1*, *S1PR1* and *TEK*, are involved in angiogenesis and cell migration, while the TG *VWF* is involved in cell adhesion. These biological processes are all closely related to cancer metastasis, indicating that Module 2 in Fig. 4A is very likely to regulate the metastasis and further development of breast cancer, which is a very worthy issue for further study.

**Fig. 3:**
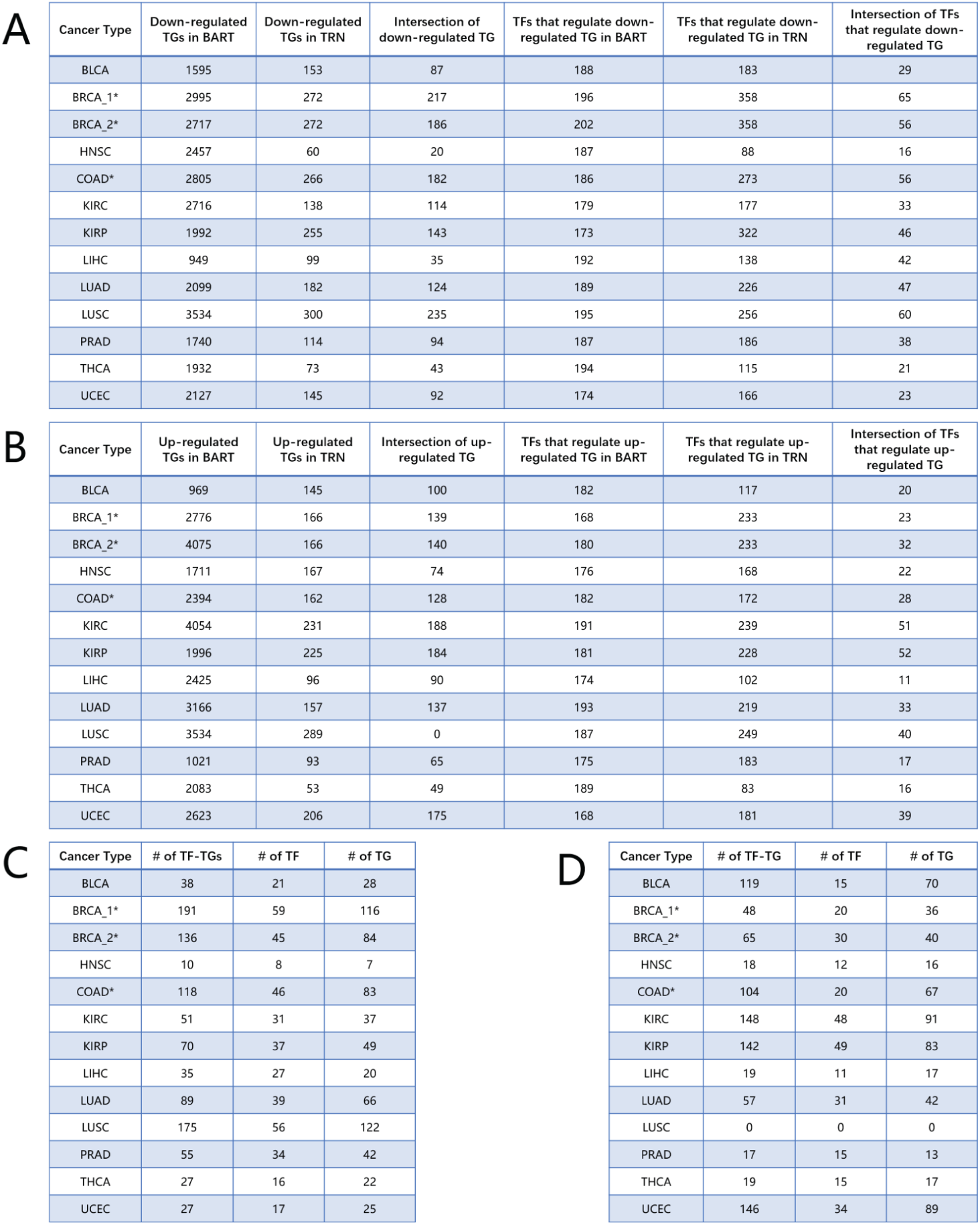
Comparison with BART Cancer. A/B: The numbers of intersections of significantly underexpressed/overexpressed TGs and their TFs. C/D: The numbers of TF-TG regulatory relationships found in the intersections of significantly underexpressed/overexpressed TGs. Note: TRN refers to the transcriptional regulatory network constructed in this study; BRCA_1*: Data is from BART Cancer’s BRCA_1 (luminal breast cancer); BRCA_2*: Data is from BART Cancer’s BRCA_2 (basal breast cancer); COAD*: Data is from BART Cancer’s COAD READ.

**Fig. 4:**
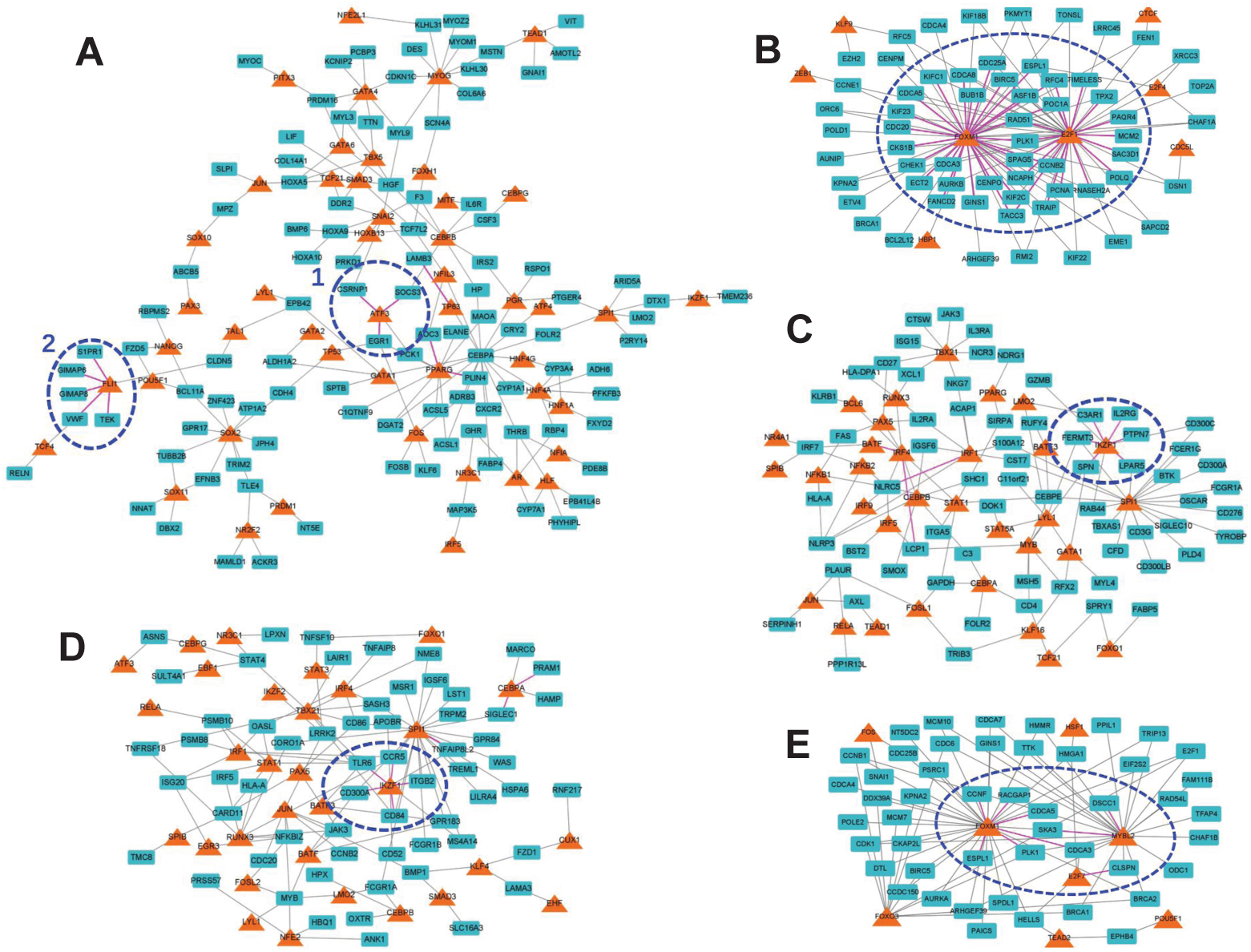
Regulatory subnetworks and important modules validated by BART Cancer and Cistrome Cancer. A: The regulatory subnetwork of underexprssed TGs in BRCA. B: The regulatory subnetwork of overexpressed TGs in BLCA. C and D: The regulatory subnetworks of overexpressed TGs in KIRC and KIRP, respectively. E: The regulatory subnetwork of overexpressed TGs in COAD. Some regulatory modules are indicated by dotted lines.

BLCA exhibited a *FOXM1/E2F1* hub module with these two TFs interacting and co-regulating targets (Fig. 4B). Studies have shown that *FOXM1* is a highly prognostic marker for non-muscle-invasive bladder cancer [42]; And *FOXM1* participated in regulating cell cycle processes and promoting the progression of bladder cancer, which makes it a potential target for the cancer treatment [43]. In addition, Logotheti *et al*. discovered that *E2F1* can promote metabolic reprogramming in bladder cancer to obtain a mixed cell phenotype to the invasiveness of the cancer [44]; Mun et al. demonstrated that *E2F1* can serve as a diagnostic marker for bladder cancer progression and a potential therapeutic target [45]. Their regulated targets are heavily involved in cell cycle/ubiquitination (*CKS1B, AURKB, BIRC5, BUB1B, CDC20, CDCA3/5/8*), proliferation (*CHEK1, CKS1B, TACC3, GINS1, CDC25A*), DNA repair/mismatch repair (*ASF1B, FANCD2, POC1A, PCNA, POLQ, RAD51, RNASEH2A*), and apoptosis (*PLK1, ESPL1, TPX2, BIRC5, MCM2*). This fully indicates that the TFs *FOXM1* and *E2F1*, which are the core regulatory factors in the module, are closely related to the occurrence and development of bladder cancer.

In KIRC and KIRP (Fig. 4C, D), enrichment analysis for the TFs and TGs revealed shared processes (TNF response, signal transduction, proliferation/apoptosis/migration regulation, cell cycle, angiogenesis, phagocytosis) and distinctions (KIRC: DNA repair, EMT; KIRP: protein metabolism/ubiquitination, Wnt regulation). Interestingly, both regulatory subnetworks for KIRC and KIRP harbored an *IKZF1*-centric module intersecting with Cistrome, though target genes differed (Fig. 4C, D). *IKZF1* targets in both cancers participated in overlapping functions like immune response, phagocytosis, inflammatory response, and cell migration, indicating *IKZF1* employs similar regulatory mechanisms to promote kidney cancer progression in both subtypes. While COAD’s module involves three Tfs: *FOXM1, MYBL2*, and *E2F7* (Fig. 4E), and they mediates cell cycle (*CCNF, CDCA3, PLK1, SKA3*), DNA repair (*CLSPN, CDCA5*), signal transduction (*FOXM1, RACGAP1*), and apoptosis (*ESPL1*), suggesting a role in colon cancer progression.

##### Novel regulatory relationships

Many TFs regulating significantly under/overexpressed TGs in our network were missed by BART Cancer. Enrichment analysis showed these ‘complement set’ TFs regulate critical processes (cell differentiation, fate commitment, proliferation, EMT, immune processes, apoptosis) (Fig. 5A, B). Examples in BRCA include:

**Underexpressed TG regulators**: There are 297 TFs not identified by BART Cancer. For example, the relationships of MYC regulating CEBPA, and CEBPB regulating IL6R (IL-6 receptor) in our TRN were validated elsewhere [46, 47] but absent in BART/Cistrome.

**Overexpressed TG regulators**: There are 207 TFs not identified by BART Cancer. Validated relationships missed in BART/Cistrome include TP53-NUAK2 (p53 network), WT1-HOXB9 (WT1-dependent regulation), and RUNX2-IBSP (bone gene regulation).

Notably, we still identified more E2F family members (E2F2, E2F3, E2F7, E2F8) than BART Cancer. These regulate cell cycle, genome integrity, and DNA damage/apoptosis responses (Fig. 5C) [48].

In summary, our multi-omics approach delineated pan-cancer TRNs, identifying conserved and cancer-specific modules driving oncogenesis. These findings provide a foundation for targeting dysregulated TFs and pathways in precision oncology.

**Fig. 5:**
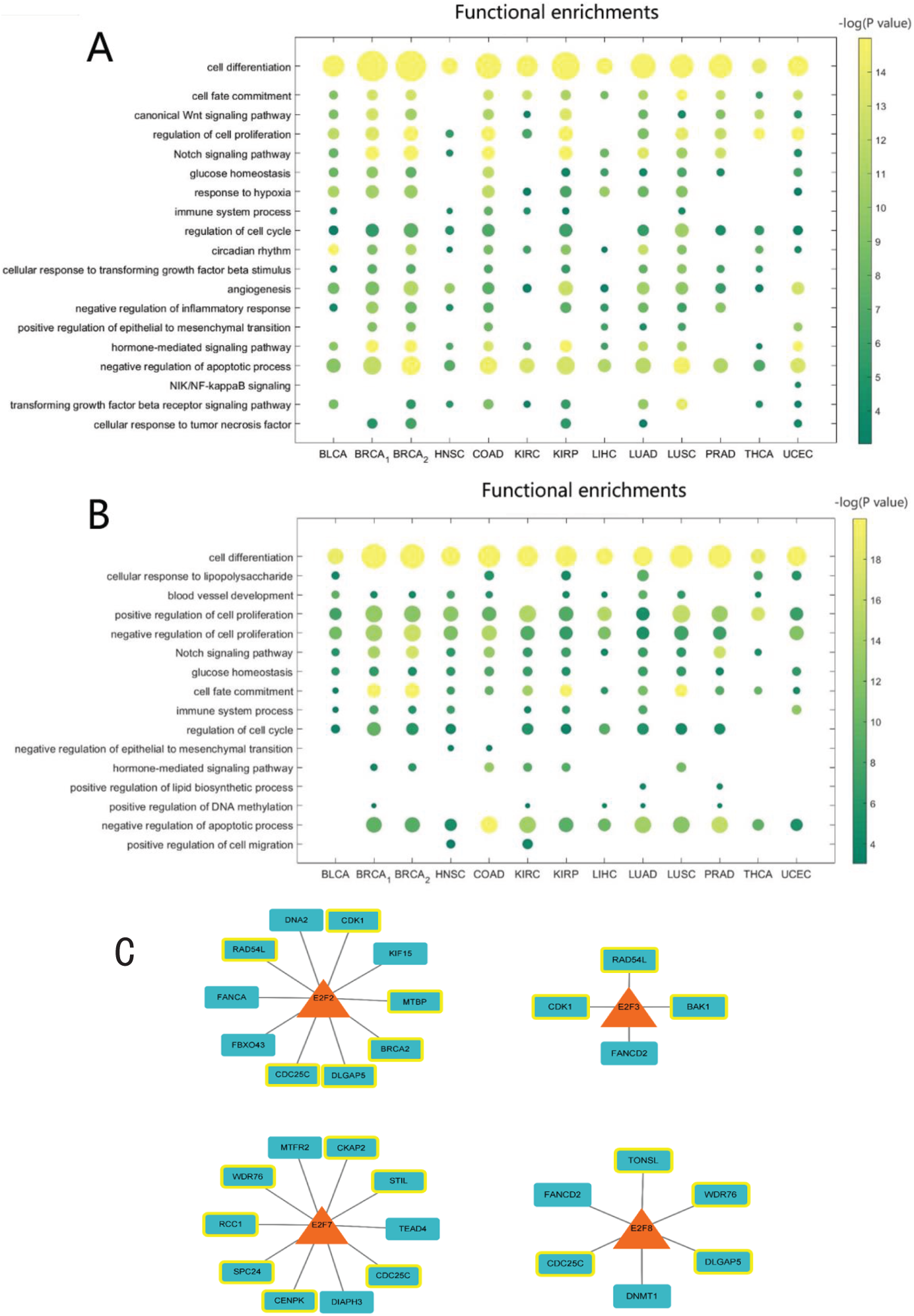
A-B: GO enrichment analysis of TFs not contained in BART Cancer (A: for TFs regulating underexpressed TGs; B: for TFs regulating overexpressed TGs). C: Target genes regulated by the *E2F* family in our constructed TRNs, however these TFs are not captured by the BART Cancer resource (The yellow box indicates that the gene is involved in cell division, DNA damage and apoptosis).

### 3.2 Cancer transcriptional regulatory networks by integrating RNA-seq data, ATAC-seq data, and DNA methylation

DNA methylation, as an important epigenetic modification, can affect the structure of chromatin and participate in gene expression regulation [7]. Methylation of DNA CpG sites in gene promoter regions is generally thought to inhibit the transcription process of genes, and one possibility is that methylated DNA prevents transcription factor binding. In view of this, when constructing regulatory networks based on RNA-seq, ATAC-seq, and DNA methylation data, we focused on the binding of transcription factor TFs to hypometylation regulatory elements.

#### 3.2.1 Pan-cancer transcriptional regulatory networks reveal a critical subnetwork involving core transcription factors

By integrating RNA-seq, ATAC-seq and DNA hypomethylation data, we constructed transcriptional regulatory networks for 15 cancer types. In order to identify important regulatory modules conserved in various cancers, we first examined the overlap of the TF-TG relationships and the overlap of TGs between any two types of cancers (Fig. 6A). Remarkably, we found that regulatory relationships recurrent in at least six cancers formed a large, interconnected subnetwork, demonstrating the possible synergistic effect that might exist among these regulations (Fig. 6B).

Transcription factor *TBP* is the largest hub gene in this connected subnetwork, which is consistent with its core role in gene transcription regulation. *TBP* is a core component of the *TFIID* complex and regulates promoter regions of a large number of coding and non-coding genes. There have been many reports indicating that *TBP* is involved in the occurrence and development of various cancers, including lung cancer [49], colorectal cancer [50], etc. The second hub gene is the transcription factor *SP2*, which is a member of the *SP* family and is widely involved in various physiological and pathological processes such as cell proliferation, differentiation, apoptosis, and embryonic development. In terms of tumorigenesis, *SP2* can directly regulate the proteins related to glioma cell formation and multiple target genes enriched in the non-small cell lung cancer pathway [52, 53], and may be a potential new therapeutic target for gastric cancer [54].

In fact, apart from the two hub genes mentioned above which play significant roles in this regulatory subnetwork, most of the other genes (including TFs and TGs) are also involved in cancer-related biological processes (e.g., *RELA, ERF, DAZAP1, MCM7*). Furthermore, we find that some regulatory relationships in this regulatory subnetwork have been confirmed by previous studies. For instance, *TBP-NFYC* was demonstrated through protein-protein interaction experiments [55], *TBP-USP10* was revealed by the fact that the deubiquitase *USP10* can antagonize a certain activity of *TBP* [56], and *E2F1-CDCA3* has been found to affect the proliferation of cancer cells in colorectal cancer [57].

All of the above analyses demonstrate that this large connected regulatory sub-network plays a significant role in the initiation and progression of cancer, warranting further in-depth investigation.

**Fig. 6:**
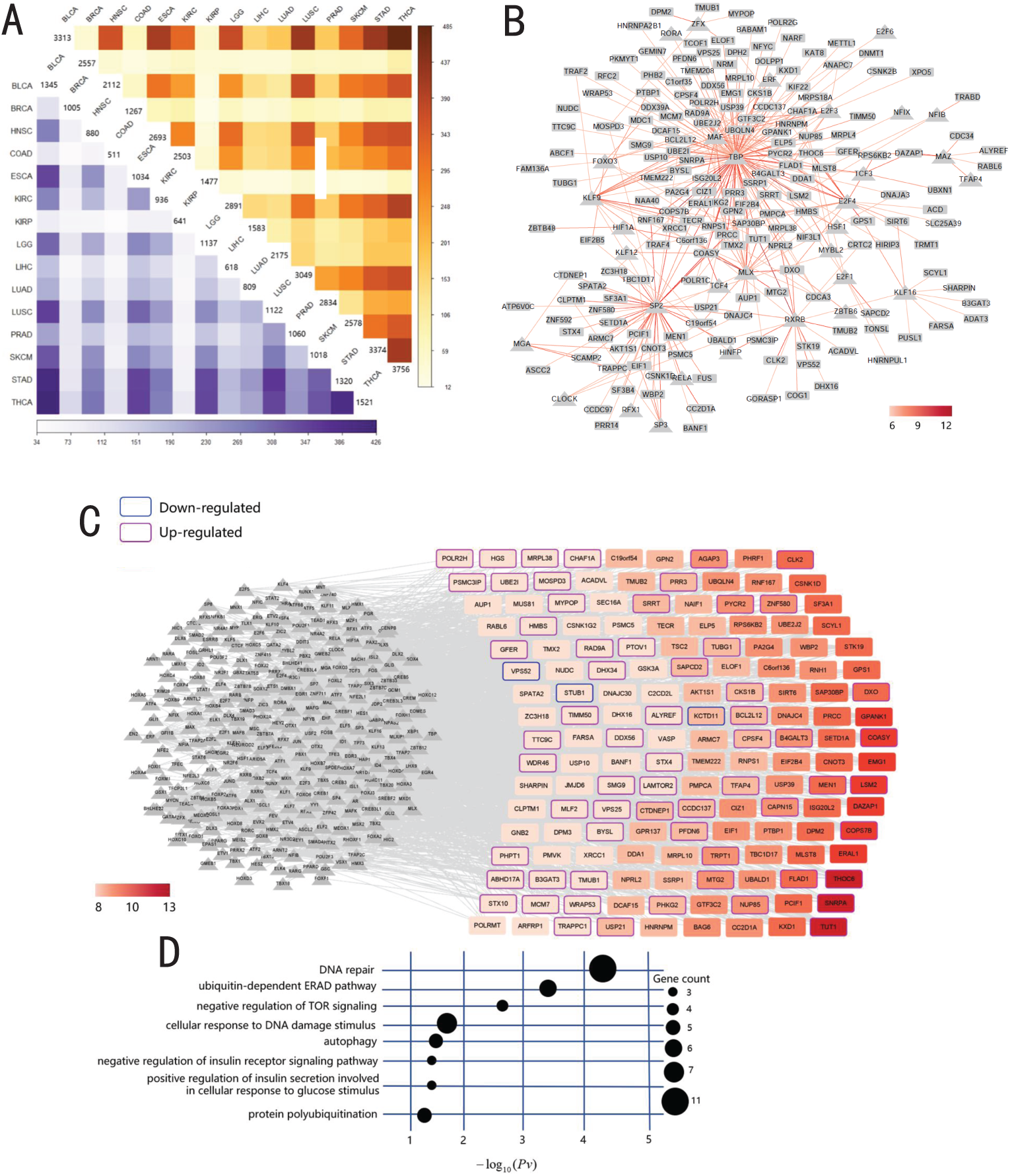
A: The overlap of the constructed TRNs between any two types of cancers. The upper and lower triangles represent the overlaps of TF-TG relationships and the TGs, respectively. B: Regulatory relationships identified in at least six types of cancer. Triangles and rectangles represent TFs and TGs, respectively. The color of the line represents the number of cancer types in which this regulatory relationship was identified. C and D: TGs appearing in at least eight cancers (C: The color of the rectangle represents the number of cancer types where the TG appears; D: Biological processes significantly enriched by the TGs).

#### 3.2.2 Recurrently dysregulated target genes in TRNs implicate core pan-cancer mechanisms

Analysis reveals numerous target genes recurrently involved in TF-TG regulatory relationships across multiple cancer types. These recurrent TGs suggest their aberrant expression is closely linked to fundamental processes driving pan-cancer development. While the regulating TFs are different and expression magnitudes of the TGs may also vary between cancers, these shared TGs converge on disrupting critical biological pathways to promote oncogenesis.

We identified TGs dysregulated in at least half (8 out of 15) of the studied cancers (Fig. 6C). Functional enrichment confirms their involvement in key cancer-related processes, such as DNA repair (e.g., *MUS81, RAD9A, STUB1, WRAP53, CHAF1A*), negative regulation of TOR signaling (e.g., *AKT1S1, NPRL2, TSC2, GSK3A*), and autophagy regulation (e.g., *USP10, SEC16A, GPR137, TBC1D17*) (Fig. 6D).

Significantly, 66 recurrent TGs exhibit differential expression, with most showing exclusive upregulation across malignancies (pink-bordered genes in Fig. 6C). This consistent overexpression pattern implies critical oncogenic functions in diverse tissue contexts.

Key overexpressed recurrent TGs and their roles include:

1) *SAPCD2*: Overexpressed in nine cancers (e.g., COAD, STAD, BRCA, LUAD, LUSC). While highly expressed during embryogenesis, *SAPCD2* is typically absent/low in healthy adult tissues. Its re-expression is linked to multiple cancers, including gastric, colorectal, lung, and breast cancer [58]. Supporting studies show high *SAPCD2* expression in colon cancer tissues versus normal tissue [59]. In breast cancer, its levels correlate with tumor size, TNM stage, and lymph node metastasis [60]. Similarly, *SAPCD2* is significantly upregulated in LUAD/LUSC and associated with poor tumor differentiation [61].
2) *MCM7* : Upregulated in eight cancers (e.g., KIRC, LUAD, LIHC, KIRP, LUSC). This conserved DNA-binding protein, part of the *MCM* family, is implicated in tumorigenesis and is a potential biomarker for malignancies [62]. In NSCLC, *MCM7* overexpression drives colony formation, proliferation, G1-S transition, and apoptosis resistance. Critically, its expression positively correlates with tumor grade and TNM stage [63]. Strong association with proliferation marker *Ki67* further confirms its role in uncontrolled cell division [64].
3) *WRAP53* : Overexpressed in KIRP and HNSC. Functioning as the antisense transcript to tumor suppressor *TP53*, *WRAP53* exhibits oncogenic properties and is a promising therapeutic target for HNSC [65]. Its overexpression predicts poor head and neck cancer prognosis [66]. Moreover, *WRAP53* is an independent prognostic factor in rectal cancer, driver of ESCA progression, and regulator of NSCLC cell behavior and radiotherapy response [67].

The recurrent dysregulation, particularly consistent overexpression, of specific TGs like *SAPCD2, MCM7*, and *WRAP53* across diverse cancers strongly suggests their involvement in universal mechanisms driving carcinogenesis. Mechanistically, they drive core cancer hallmarks: developmental reprogramming and metastasis (*SAPCD2*), sustained proliferative signaling (*MCM7*), and genome instability and therapy resistance (*WRAP53*).

Therefore, studying recurrently dysregulated TGs across various cancers is highly significant. These genes represent potential central nodes in pan-cancer networks and could serve as valuable diagnostic/prognostic biomarkers or novel therapeutic targets applicable to a broad spectrum of malignancies. Understanding their precise roles remains crucial for uncovering fundamental cancer biology and developing effective, broadly applicable therapies.

#### 3.2.3 Prognostic models identify high-risk target genes in TRNs

In cancer research, target genes with low methylation levels in their promoter regions often exhibit high expression. These TGs are of particular interest due to their potential association with cancer progression. In this study, we took two cancer types, COAD and KIRP, as examples to investigate the prognostic relevance of highly expressed TGs, their role in risk assessment, and their impact on patient survival.

In COAD, we identified *TAS2R14, FBXL13, DOCK6*, and *ADAMTS14* as candidate prognostic TGs in COAD by univariate Cox and LASSO-Cox regression analyses (Eqs. 4 and 5) (Fig. 7B, C). Among these, *TAS2R14* has been extensively reported to correlate with patient survival in multiple cancers (e.g., thyroid carcinoma, prostate cancer, ovarian cancer, breast cancer) [68, 69]. *FBXL13*, though not previously linked to prognosis, has been identified as a regulator of microtubule nucleation, centrosome homeostasis, and cell migration, suggesting its potential role in tumor progression [70]. To validate these findings, we stratified patient samples into high-risk and low-risk groups based on risk scores. We noticed that the high-risk group exhibited a higher mortality rate (Fig. 7D). Survival analysis further confirmed a significant difference in survival outcomes between the two groups (*p <* 0.05), underscoring the prognostic value of these TGs in COAD (Fig. 7E).

Analysis of KIRP identified *ANLN* as a key prognostic candidate TG. In fact, *ANLN* does have a strong correlation with cancer. In various types of tumors, overexpression of *ANLN* is associated with cell proliferation and migration, tumor mutation burden, microsatellite instability, phosphorylation level and increased immune cell infiltration, and can serve as a biomarker for pan-cancer prognosis [71]. Crucially, *ANLN* serves as a prognostic marker in numerous cancers, including colorectal cancer (associated with invasion and poorer survival) [72], pancreatic cancer (promoting progression) [73], bladder cancer (independent predictor) [74], esophageal squamous cell carcinoma (promoting proliferation, poor prognosis) [75], etc. Here in KIRP, risk stratification based on *ANLN* expression similarly showed high-risk patients experienced significantly more deaths and shorter survival time. These results highlight *ANLN*’s potential as a biomarker for KIRP prognosis.

In summary, by prognostic modeling and survival analysis based on the differentially expressed TGs in TRNs, we were able to identify some TGs associated with poor prognosis of cancer. The study of these TGs is beneficial for cancer treatment and may provide opportunities for prolonging patients’ survival time.

**Fig. 7:**
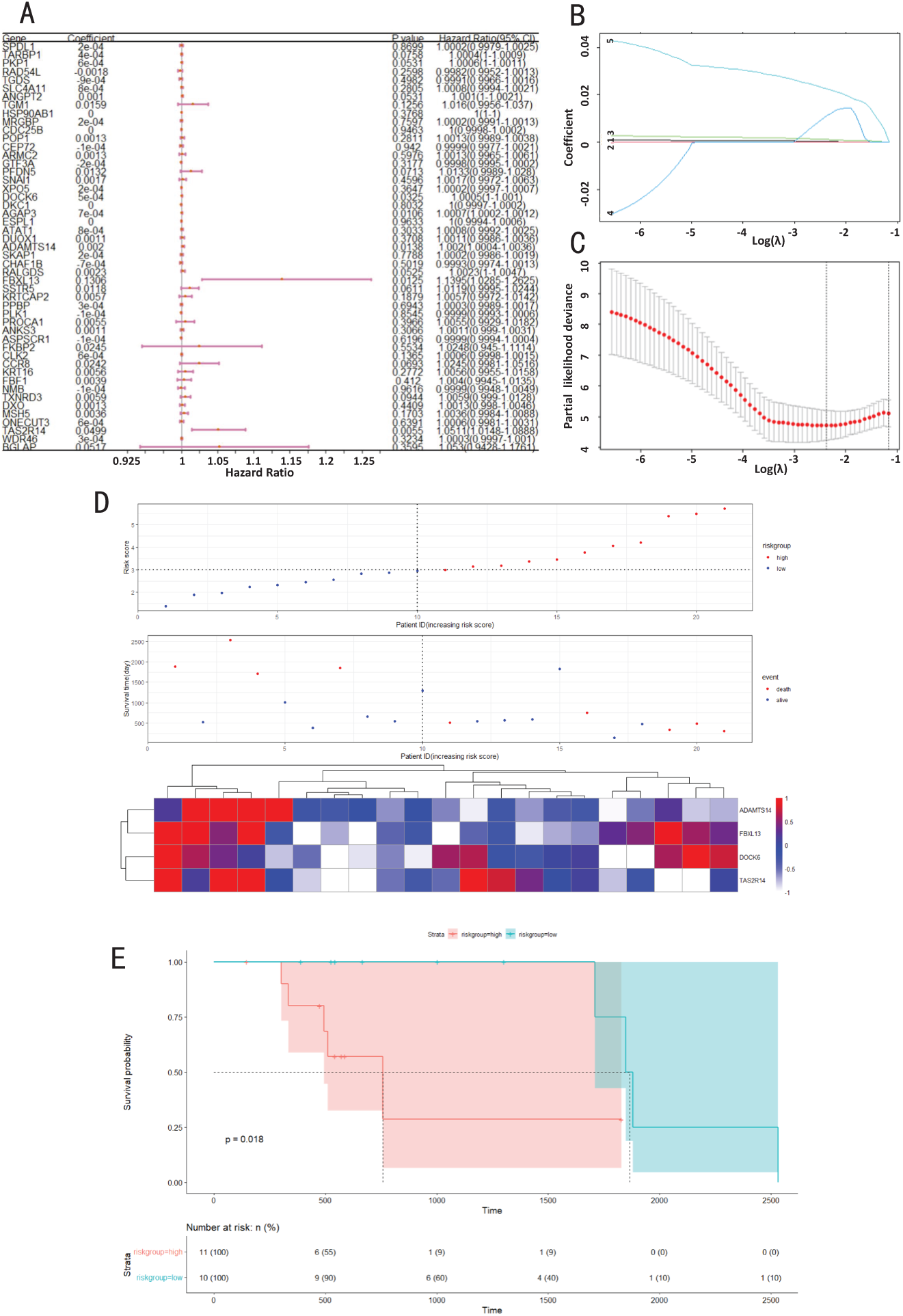
Prognostic analysis, risk assessment, and survival analysis of COAD. A: Univariate COX regression analysis. Orange dots and pink line segments denote hazard ratios and 95% confidence intervals, respectively. B: The change in the coefficient for each gene when *λ* changes in the LASSO-COX regression. Digits 1-5 represent five genes (*DOCK6, AGAP3, ADAMTS14, FBXL13* and *TAS2R14*, respectively). C: The selection of *λ* that minimizes model error in LASSO-COX regression analysis. D: Risk assessment of candidate TGs. E: Survival analysis.

#### 3.2.4 Enhancer-associated regulatory relationships underlie cancer pathogenesis

Enhancer methylation plays a crucial role in cancer. There were studies demonstrating that DNA methylation of enhancers is more closely related to the expression of cancer-related genes than promoter DNA methylation [76]. Changes in enhancer methylation can indicate patient prognosis and may promote cancer progression through cancer cell plasticity [77].

Here we intend to investigate regulatory effects of enhancer methylation in pan-cancer. Considering we have integrated DNA methylation in the model, we examine enhancers that intersect with the REs in the TRNs. First, we collected enhancers related to various cancer types from the HEDD database [36], in which we matched the cancer types in this study and counted the intersected enhancers with our REs (Supplementary Material 1, Table S2). Except for HNSC, ESCA, KIRP, LUAD, and SKCM, the REs of other cancers were all found to be more or less involved in cancer-related enhancers. Further, we examined the TF-RE-TGs regulatory relationships where enhancers are involved in. In fact, the HEDD database also contains the information for the TFs that bind to enhancers and the TGs they regulate. We focused on regulatory relationships where the target genes were consistent with those in the HEDD. Such integration of cancer-related enhancers with TRNs revealed critical TF-RE-TG interactions.

In BLCA, the *TBP-WRAP53* regulatory relationship (located at enhancer EnhancerID:328441) was predicted by both our TRN and the HEDD database (Fig. 8A). As stated above, transcription factor *TBP* contributes to cancer development. *WRAP53* is also highly expressed and serves as a prognostic marker in multiple cancers [78], suggesting the *TBP-WRAP53* axis influences bladder cancer progression. Our TRN additionally shows *KLF9* regulates *WRAP53* (Fig. 8A). *KLF9* targets genes that suppress bladder cancer proliferation and inhibit progression [79], and modulates cancer cell proliferation/differentiation in various cancers including lung, breast, gastric, prostate, colon, endometrial, and liver cancers [80]. This implies the *KLF9-WRAP53* relationship critically impacts bladder cancer proliferation and migration.

For BRCA (Fig. 8B), RPS6KB2 – a known mTOR downstream target involved in protein translation, cell growth, and proliferation – is extensively studied in cancer. Sridharan *et al*. highlighted its significant role in regulating breast cancer cell death [81]. The HEDD database predicted *TCF12* binding to the *RPS6KB2* enhancer (ID:246437), while our TRN shows regulation by *TCF4*. Given that *TCF4* and *TCF12* are homologous genes with typically similar functions and regulatory pathways, the *TCF4-RPS6KB2* and *TCF12-RPS6KB2* relationships likely play analogous roles in breast cancer. *TCF4* exhibits diverse cancer roles, including tumor suppression in breast cancer [82]. Additionally, *KLF1* binds a breast cancer-related enhancer (ID:1764767) on *RPS6KB2*. While no direct breast cancer link for *KLF1* is established, its importance in other cancers (such as non-small-cell lung cancer, cervical cancer, and gastric cancer) [83] warrants further investigation of *KLF1-RPS6KB2* in breast cancer.

In LIHC, HEDD database analysis shows enhancers 460287 and 460291 (largely overlapping RE1: chr20:50112947C50113448 and RE2: chr20:50153574C50154075) regulate *CEBPB* (Fig. 8C). The target gene *CEBPB* exhibits critical roles across cancers: it activates drug resistance in lung/nasopharyngeal cancers [84], promotes head and neck squamous cell carcinoma, breast/ovarian cancers [85], suppresses growth/metastasis in colorectal/breast/cervical tumors [86], correlates with favorable prognosis in metastatic melanoma [87], and inhibits osteosarcoma proliferation [88]. Given *CEBPB* s broad oncogenic relevance, the regulatory axes *HES2*-RE1-*CEBPB*, *HES2*-RE2-*CEBPB*, and *TFEC*-RE2-*CEBPB* are biologically significant. Furthermore, transcription factor *TFEC*, as a member of the MiT family, is considered an oncogene, and dysregulation of this family can lead to melanoma, renal cell carcinoma, and some sarcomas [89]; and transcription factor *HES2* is upregulated in BMP9-induced ovarian cancer cells, pancreatic cancer, mismatch repair-deficient colon cancer, cervical cancer, and head and neck tumors [90]. All these further support the above three regulatory relationships as key research targets in liver cancer.

Regarding THCA, among the 73 enhancer-containing regulatory relationships (Supplementary Material 1, Table S2), we identified a module centered on transcription factor *SP2* (Fig. 8D). *SP2* regulates three target genes (*C6orf136, ABCF1, PRRT1*), all implicated in cancer development [91]. *TBP* co-regulates *C6orf136* and *ABCF1* with *SP2* ; both TFs are cancer-associated (Fig. 6B). *ETS1*, also linked to poor cancer survival [92], regulates *C6orf136*. Among the other four TFs (*E2F3, NFIB, KLF12, SOX11*) regulating *ABCF1*, *E2F3*’s close relationship with cancer has been displayed in Fig. 5C; *NFIB* and *KLF12* have been shown to play important roles in cancer growth, proliferation, migration, and prognosis [93, 94]; and *SOX11* has been demonstrated to promote cancer progression in various cancers by regulating cells or stem cells [95]. On the other hand, another TF regulating *PRRT1* in the module is *GATA2*, which plays a critical role in regulating gene transcription involved in hematopoietic and endocrine cell lineage development and proliferation, and promotes cancer invasiveness, proliferation, and tumorigenicity in various cancers [96]. Collectively, the cancer-related roles of these interconnected TFs and targets strongly suggest this *SP2*-centered module plays a crucial role in THCA development, warranting further study.

Analysis of regulatory relationships across multiple cancers reveals recurring interactions, suggesting pan-cancer significance.

Enhancer 1764767 regulates *RPS6KB2* and is predominantly contained within REs in BRCA, LIHC, and THCA (Fig. 8E). The *NFIB-RPS6KB2* relationship occurs in all three cancers, highlighting its importance. Both genes are established in cancer development [81, 93]. The *MAZ-RPS6KB2* relationship is predicted in BRCA and THCA. Transcription factor *MAZ* exhibits context-dependent roles: it activates some oncogenes while inhibiting others. *MAZ* has dual functions in breast cancer (inhibiting invasiveness yet promoting proliferation [97]), influences the tumor microenvironment and immune escape in liver cancer [98], promotes prostate cancer metastasis [99], inhibits migration/invasion in gastric/pancreatic cancers [100], and is highly expressed in colon cancer inflammation [101]. This implicates both *MAZ-RPS6KB2* and *NFIB-RPS6KB2* in regulating cancer growth, invasion, and metastasis.

**Fig. 8:**
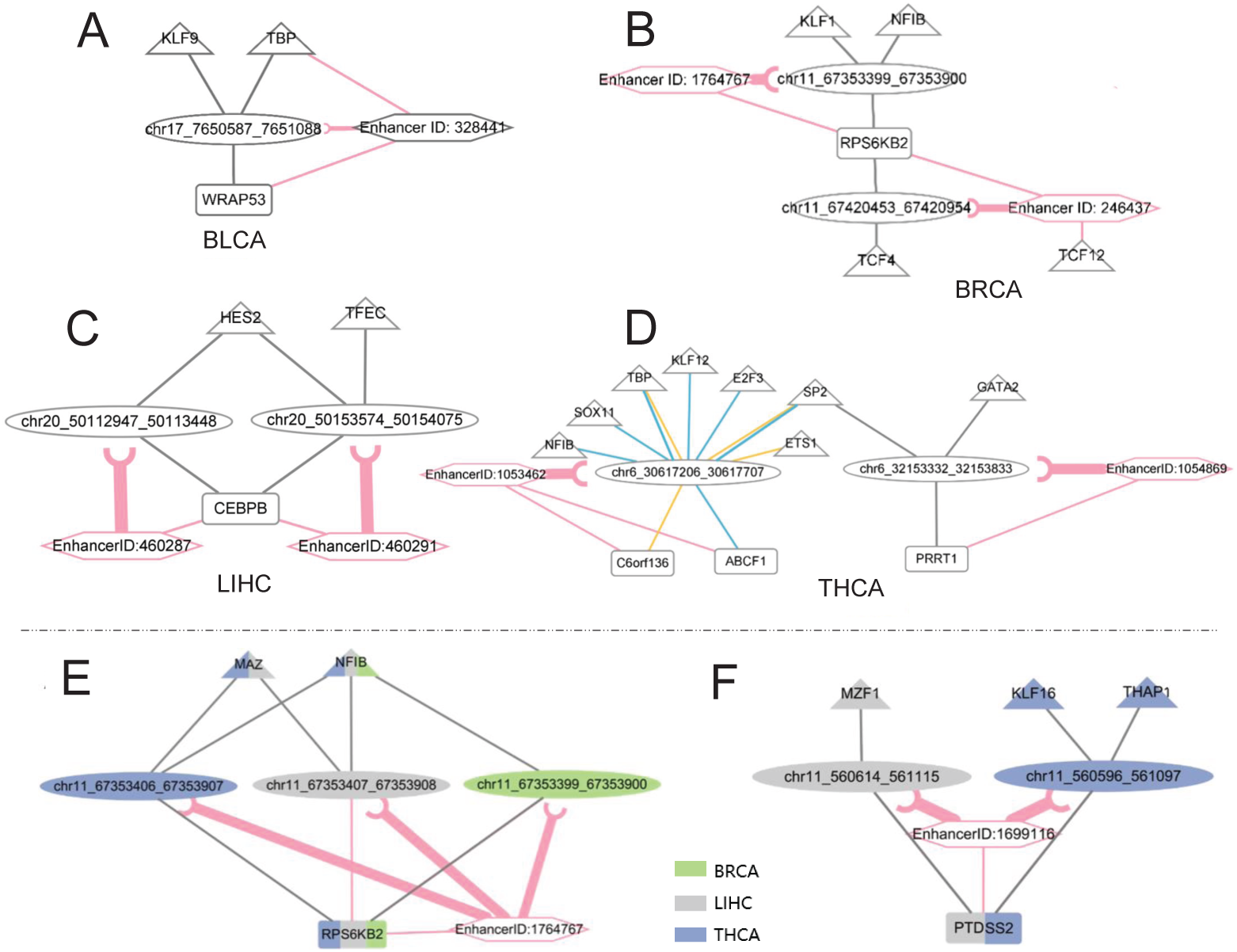
Multiple regulatory relationships TF-RE(-enhancer)-TGs in cancers. Among them, the lines in gray and pink represent regulatory relationships in the constructed TRN and in the HEDD database, respectively. The thickness of the line between REs and enhancers represent the overlap degree of REs and enhancers. A: BLCA; B: BRCA; C: LIHC; D: THCA; and multiple regulatory relationships involved in multiple cancer types (E: BRCA, LIHC and THCA; F: LIHC and THCA).

Enhancer 1699116 co-regulates *PTDSS2* in LIHC and THCA, fully encompassed by the respective regulatory elements (Fig. 8F). *PTDSS2* functions in signaling, coagulation, and apoptosis, and its overexpression accelerates metastasis [102]. In LIHC, the oncogenic transcription factor *MZF1* regulates *PTDSS2*, aligning with its pro-metastatic role. In THCA, *PTDSS2* is regulated by *KLF16* and *THAP1*. *KLF16* generally exhibits anti-cancer activity, inhibiting growth/migration in multiple cancers [103, 104]. *THAP1* regulates endothelial proliferation and the G1/S transition and shows differential expression in papillary thyroid cancer [105], suggesting pathogenic involvement. The regulatory axes *MZF1-PTDSS2, KLF16-PTDSS2*, and *THAP1-PTDSS2* thus appear significant in pan-cancer.

#### 3.2.5 DNA hypomethylation at CpG sites modulates oncogene regulation

The integration of DNA methylation provides more information for the understanding of the transcriptional regulation. Here we mainly investigate the role of methylation at CpG sites (mCpG) within REs controlling the oncogenes. For example, as the oncogenes associated with promoter hypomethylation [23], *FKBPL, BTK*, and *PRRT1* are also most prevalent in our pan-cancer TRNs (appearing in five cancers). The TFs regulating them participate in key cance-related processes (Supplementary Material 1, Fig. S5). For instance, *BTK*-regulating TFs are involved in protein ubiquitination and hypoxia response; *FKBPL*-regulating TFs in cell differentiation and Notch pathway inhibition; and *PRRT1*-regulating TFs in glucose homeostasis, angiogenesis, and endothelial cell migration.

*BTK*, a membrane protein driving malignant B-cell proliferation, is crucial for B-cell function and a clinical cancer treatment target. In HNSC, *BTK* methylation affects its isoforms (*BTK-p80/p65*), promoting cancer [106]. We identified five distinct REs regulating *BTK* across BRCA, ESCA, KIRC, PRAD, and SKCM (Fig. 9A). In PRAD, RE chrX 101348503 101349004 contains CpG site cg06818142; upstream TF *MYB* binding forms the *MYB-BTK* regulatory axis. *MYB*’s established link to prostate cancer involves androgen receptor interaction, where its overexpression increases risk [107]. *MYB* (*c-Myb*) also regulates hematopoiesis and is aberrantly expressed in leukemia, lymphoma, and solid tumors (breast, ovarian) [108], acting as an oncogenic inducer promoting growth and metastasis in liver, colon and breast cancer [109].

Notably, REs frequently overlap between cancers, especially when bound by the same TF. For example, ESCA and SKCM REs overlap significantly (one base difference), bound by *NPAS1* and *NFE2*, respectively (and both bound by *NPAS2*). *NPAS2* influences cancer via cell cycle regulation and DNA damage response, while *EPAS1* regulates cancer prognosis and drug resistance [53]. Similarly, KIRC and BRCA REs overlap (one base difference), both bound by *NFE2*, plus *HIF1A, HOXB4*, and *TFEB, TFEC, SREBF2*, respectively. *HIF1A* is implicated in various cancers [110]; *TFEB/TFEC* regulate lysosome biogenesis, energy homeostasis, and autophagy [89]; and *SREBF2* promotes migration and poor prognosis in breast, lung, and ovarian cancers [111]. Integrating CpG sites revealed many within overlapping REs affecting multiple cancers (e.g., cg06162422 in KIRC/ESCA overlap; cg18502619 and cg19528502 in BRCA/ESCA/KIRC/SKCM overlap), influencing *BTK* methylation levels.

*FKBPL*, involved in immune regulation, protein folding/transport, and cell cycle control, is cancer-relevant. In PRAD, only one RE was found, which was bound by *ATF1, ELK3, ELF5*, and *TBP* (Fig. 9B). *ELF5*, expressed in *AR*-activated prostate cancer cells, binds *AR* to negatively regulate its activity, inhibiting EMT and metastasis [112]. In KIRC, five REs were identified, two of which (chr6 32178631 32179132, chr6 32127594 32128095) were bound by KIRC-specific TFs (*DLX1, DLX2, HOXB4, MYB, MYBL2*). Three others overlap with STAD REs, bound by TFs like *MAZ* and *KLF7* (present in STAD/KIRC). *KLF7*inhibits gastric cancer cell proliferation/invasion/migration and is a survival predictor [113]. In a word, CpG sites within overlapping regulatory areas affect multiple cancers, and KIRC-specific TFs (*DLX1, DLX2, HOXB4, MYB, MYBL2*) bind cancer-specific REs and CpG sites (Fig. 9B).

REs regulating *PRRT1* differ substantially across LGG, LIHC, SKCM, PRAD, and THCA (Supplementary Material 1, Fig. S6). RE chr6 31958823 31959324 appears in LGG and SKCM, bound by *ETV2* (LGG) and *ETS1* (both). *ETV2* amplification occurs in cancers and is essential for tumor angiogenesis [114]. In PRAD, we only found one RE that regulates *FKBPL*. This RE was bound by *EGR1, FOXL2, ETV4*, and *NFE2*. These TFs also only appeared in PRAD. In LIHC, we identified nine REs. Among them, four did not overlap with any other REs, and they were bound to the TFs (*FOXA3, FOXD2, MLXIPL, PITX1, CTCFL*) that are only present in LIHC. In short, similar as the situation in *PRRT1*, CpG sites in overlapping areas affect various cancers, and PRAD-specific TFs (*EGR1, FOXL2, ETV4, NFE2*) and LIHC-specific TFs (*FOXA3, FOXD2, MLXIPL, PTX1, CTCFL*) regulate *PRRT1* via cancer-specific REs and CpG sites (Supplementary Material 1, Fig. S6).

**Fig. 9:**
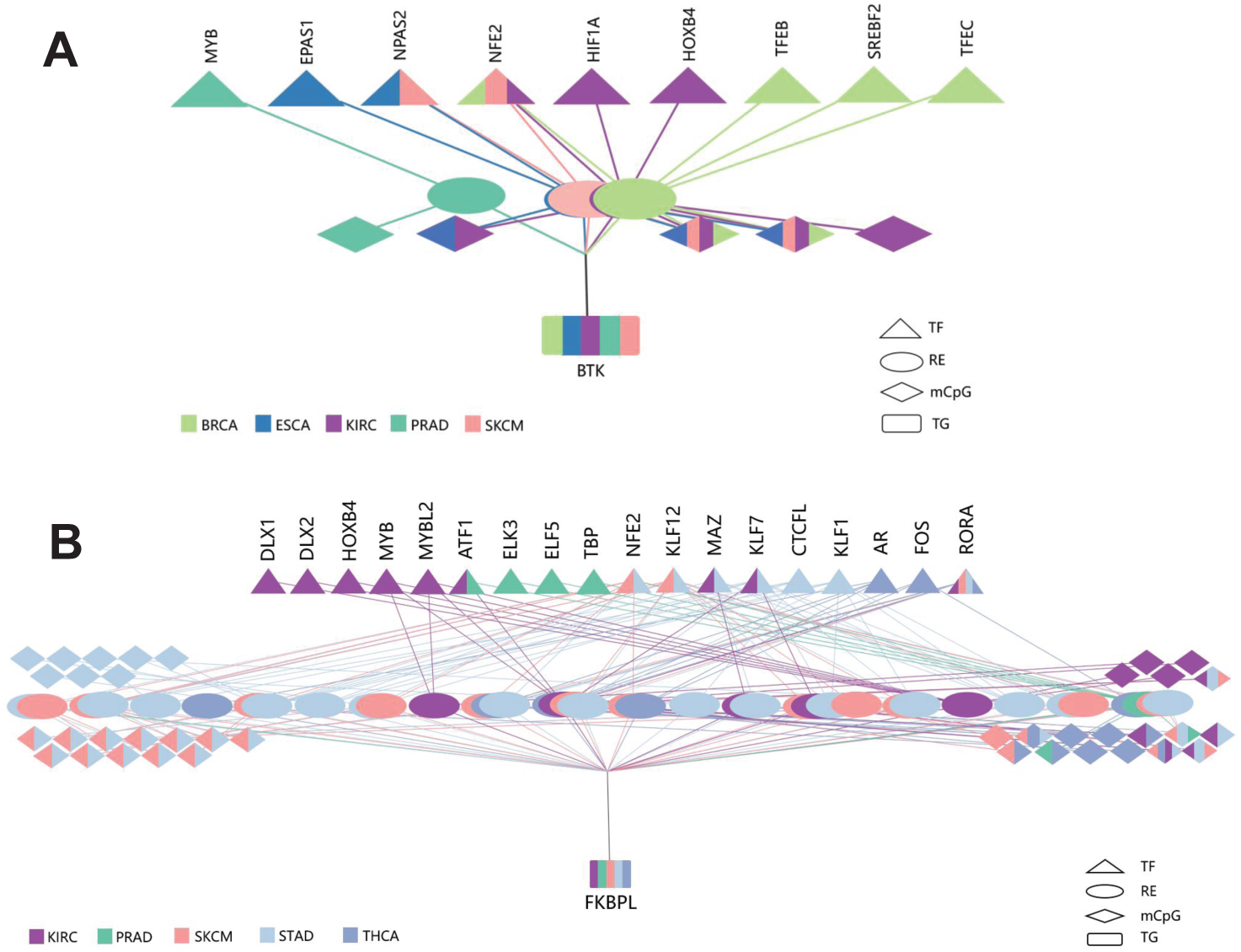
Multiplex complex regulatory relationships that regulates TGs in multiple cancer types. A: TF-RE(-mCpG)-*BTK* in BRCA, ESCA, KIRC, PRAD, and SKCM. B: TF-RE(-mCpG)-*FKBPL* in KIRC, PRAD, SKCM, STAD and THCA. Here mCpG represents the methylated CpG site on DNA.

Analysis of these three methylation-affected oncogenes demonstrates that DNA methylation significantly influences transcriptional regulation, potentially altering target gene states to induce or inhibit cancer cell growth, proliferation, and metastasis, warranting further investigation.

To sum up, our multi-omics approach delineated a pan-cancer TRN enriched with conserved regulatory modules, prognostic TGs, and enhancer-driven interactions. DNA hypomethylation at CpG sites further elucidated mechanisms underlying oncogene activation. These results provide a framework for targeting shared regulatory vulnerabilities across cancers.

## 4 Discussion

This study advances our understanding of pan-cancer transcriptional regulation by integrating multi-omics data to reconstruct context-specific transcriptional regulatory networks. By leveraging RNA-seq, ATAC-seq, and PPI or DNA methylation profiles across 16 TCGA cancer types, we systematically decoded the interplay between TFs, REs, and TGs, revealing conserved and cancer-specific mechanisms driving tumorigenesis.

A key strength of this work lies in its integrative framework. Extending the PECA model to incorporate PPI networks or methylation data improved the biological interpretation of inferred TF-RE-TG relationships. For instance, hypomethylated REs (*β <* 0.3) were prioritized to refine regulatory interactions, uncovering methylation-sensitive oncogenes such as *BTK* and *PRRT1*. The identification of recurrent regulatory modules (e.g., *TBP/SP2*-centered subnetworks) across multiple cancers underscores the existence of shared transcriptional vulnerabilities. These modules, enriched in apoptosis, cell differentiation, and DNA repair pathways, align with known oncogenic processes but also reveal novel interactions (e.g., *MYC-CEBPA*) absent in established databases like BART Cancer [27] and Cistrome Cancer [28]. Such findings highlight the value of multi-omics integration in capturing the complexity of cancer regulatory landscapes.

The study further elucidates the role of enhancers and DNA methylation in transcriptional dysregulation. Enhancer-associated interactions, such as *TBP-WRAP53* in BLCA and *SP2-ABCF1/PRRT1* in THCA, demonstrate how spatial chromatin organization cooperates with TFs to activate oncogenic programs. DNA hypomethylation at specific CpG sites (e.g., cg06818142 in PRAD) was shown to modulate TF binding and TG expression, providing mechanistic insights into epigenetic-driven tumorigenesis. These results bridge the gap between epigenetic modifications and transcriptional control, offering a framework for targeting methylation-sensitive regulatory nodes.

Prognostic models derived from TRN analysis identified high-risk TGs (e.g., *TAS2R14* in COAD, *ANLN* in KIRP) with significant survival implications. The integration of LASSO-Cox regression and Kaplan-Meier analysis validated their clinical relevance, emphasizing the translational potential of TRN-derived biomarkers. However, while these models demonstrate promise, their generalizability requires validation in independent cohorts, particularly given the heterogeneity of tumor microenvironments across cancer types.

## 5 Conclusion

In conclusion, this study provides a pan-cancer atlas of transcriptional regulatory networks, unifying multi-omics data to decode the interplay between genetic and epigenetic drivers of cancer. Key contributions include: 1) identification of conserved regulatory modules (e.g., *TBP/SP2* hubs) and cancer-specific subnetworks driving oncogenic processes; 2) mechanistic insights into enhancer-mediated and methylation-sensitive transcriptional dysregulation; 3) prognostic biomarkers (e.g., *TAS2R14, ANLN*) with potential clinical utility in risk stratification; and 4) novel TF-TG interactions (e.g., *MYC-CEBPA*) that expand existing regulatory databases.

These findings underscore the power of integrative multi-omics approaches in precision oncology. By mapping regulatory vulnerabilities, this work lays the groundwork for developing therapies targeting master regulators or epigenetic modifiers to disrupt oncogenic networks. Future efforts should focus on experimental validation and translational applications to harness these insights for improved patient outcomes.

## Acknowledgements

We would like to thank Dr. Kangning Dong and Dr. Zhanying Feng for the suggestions and the kind help. The computations were partly done on the high performance computers of State Key Laboratory of Scientific and Engineering Computing, Chinese Academy of Sciences.

## Author contributions

J.Z. conceived and supervised the study. Y.Y. gathered the data and performed the methods. J.Z. and Y.Y. interpreted the results. All authors wrote, revised and approved the manuscript.

## Data availability

All biological data used in this study are available on public data platforms. RNA-seq was download from TCGA database (https://portal.gdc.cancer.gov/). ATAC-seq was from the chromatin accessibility atlas [11]. TF motif information was integrated from the NCBI [30], UCSC Genome Browser [31], and JASPAR [32] databases. DNA methylation data was obtained from the TCGA Infinium HumanMethylation450k platform. The PPI network was from STRING [33].

The code for PEKA model was from [19]. Any additional information that supports the findings of this study is available from the corresponding author upon reasonable request.

## Declarations

### Ethics approval and consent to participate

Not applicable.

### Consent for publication

Not applicable.

