## Supplemental Tables S1-S2 and Figures S1-S6 for "Pan-cancer transcriptional regulatory network analysis reveals key drivers and epigenetic modulators in tumorigenesis"

### Cancer types used in this study

The pan-cancer multi-omics data used in this study (including gene expression, chromatin accessibility, and DNA methylation) are from The Cancer Genome Atlas (TCGA). The cancer types and their abbreviations are listed in Table S1.

**Table S1** Cancer types used in this etudy and their abbreviations

| Cancer type | Abbreviation |
|---|---|
| bladder urothelial carcinoma | BLCA |
| invasive breast carcinoma | BRCA |
| colon adenocarcinoma | COAD |
| esophageal carcinoma | ESCA |
| head and neck squamous cell carcinoma | HNSC |
| kidney renal clear cell carcinoma | KIRC |
| kidney renal papillary cell carcinoma | KIRP |
| lower grade glioma | LGG |
| liver hepatocellular carcinoma | LIHC |
| lung adenocarcinoma | LUAD |
| lung squamous cell carcinoma | LUSC |
| prostate adenocarcinoma | PRAD |
| skin cutaneous melanoma | SKCM |
| stomach adenocarcinoma | STAD |
| thyroid carcinoma | THCA |
| uterine corpus endometrial carcinoma | UCEC |

### The supplementary figures and tables for the Results

First, the 'Note' for Fig. 2A-D in the main text is as follows:

Note: (positive) regulation of cell proliferation[1]: including positive regulation of epithelial cell proliferation, positive regulation of stem cell proliferation, and regulation of cell proliferation; negative regulation of cell proliferation[2]including negative regulation of keratinocyte proliferation, and regulation of cell proliferation(negative regulation of) angiogenesis[3]: where *HOXA5, KLF4* and *PPARG* are involved in the negative regulation of angiogenesis; (positive regulation of) apoptotic process[4]: *NR3C1* and *TP63* are related to apoptotic process, and others are involved in the positive regulation of apoptosis; positive regulation of cell proliferation[5]: including positive regulations of cell proliferation, endothelial cell proliferation, glial cell proliferation, epithelial cell proliferation, and stem cell proliferation.

Next, this section includes the Figs. S1-S6 and Table S2, which have been cited in the **Results** of the main text.

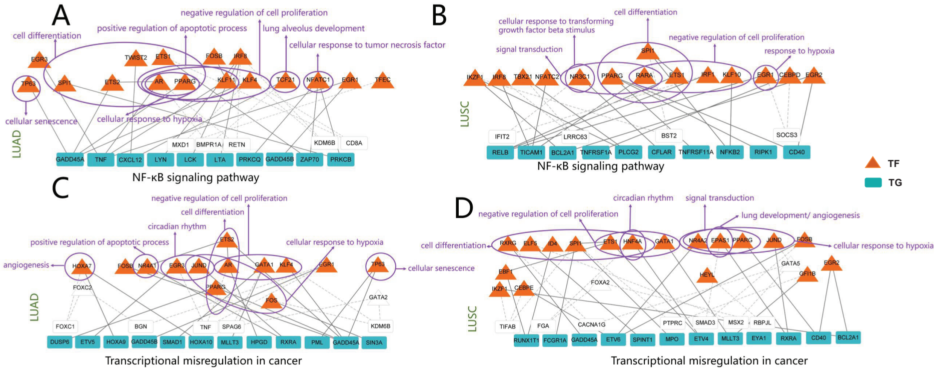


**Fig. S1** Function and regulatory subnetworks of underexpressed TFs regulating KEGG signaling pathways in LUAD and LUSC. A, B: The TGs belong to the NF-$\kappa$B signaling pathway (A: LUAD, B: LUSC). C, D: TGs belong to transcriptional misregulation in cancer (C: LUAD, D: LUSC). The white rectangles and triangles are other TGs and TFs added to make the subnet connected.

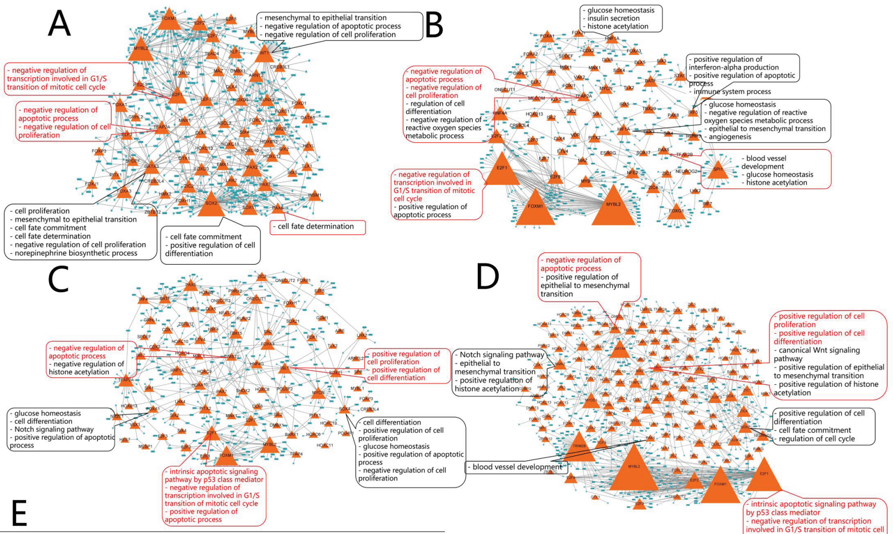


E

| GO term | -logP in BRCA | TFs in BRCA | -logP value in UCEC | TFs in UCEC |
|---|---|---|---|---|
| cell differentiation | 20.46 | FOXA1, FOXA3, DLX4, DLX5, ZIC2, NR2F6, ELF3, SPDEF, FOXD3, FOXD2, FOXD1, SOX11, SOX12, SOX2, FOXS1, MYBL1, HMX2 | 35.49 | FOXA1, FOXA3, DLX4, DLX5, ZIC2, NR2F6, ELF3, SPDEF, EHF, DLX3, SPI1, DLX6, HNF4G, FOXI1, SOX17, MECOM, HNF4A, ZIC1, SOX4, CDX2, VDR, ETV4, BATF, PAX8, ELF5, FOXA2 |
| positive regulation of cell proliferation | 5.46 | MAZ, TFAP2B, XBP1, KLF5, CDX2, E2F3, SOX9, FOXM1, SOX4 | 5.76 | MAZ,ARNT2, LEF1, SOX11, FOXM1, HOXC10, RUNX2, HMX2 |
| cell fate commitment | 3.54 | ONECUT2, ONECUT1, SOX6 | 6.84 | SOX2, GATA5, GATA3, SOX12 |
| negative regulation of apoptotic process | 4.55 | TFAP2A, ARNT2, SIX4, WT1, PAX7, LEF1, PAX2 | —— | —— |
| mesenchymal to epithelial transition | 7.74 | WT1, GATA3, PAX2 | —— | —— |
| norepinephrine biosynthetic process | 3.71 | GATA3, INSM1 | —— | —— |
| transforming growth factor beta receptor signaling pathway | —— | —— | 4.45 | SPI1, ONECUT2, FOXH1, ONECUT1 |
| immune system process | —— | —— | 3.57 | IRF7, IRF5, IRF6 |
| Notch signaling pathway | —— | —— | 6.13 | FOXA1, ONECUT1, SIX1, SOX9, TP63 |

F

| GO term | -logP in LUAD | TFs in LUAD | -logP value in LUSC | TFs in LUSC |
|---|---|---|---|---|
| Notch signaling pathway | 6.47 | ONECUT1, FOXA1, HES7, MESP1, SIX1 | 8.59 | ONECUT1, DLX1, DLX2, SNAI2, HOXD3, SOX9, TP63 |
| positive regulation of cell proliferation | 9.08 | PDX1, CDX2, SOX11, FOXM1, ISL1, HOXC10, E2F3, HSF4, POU3F2, SOX4, RUNX2 | 14.74 | PDX1, CDX2, SOX11, FOXM1, ISL1, HOXC10, E2F3, HSF4, POU3F2, TBX1, TFAP2B, RARG, LEF1, POU3F3, FOSL1, KLF5, SOX9, HMX2 |
| cell fate commitment | 3.71 | ONECUT1, ONECUT2, NR2E1 | 9.87 | ONECUT1, ONECUT2, NR2E1, SOX2, GATA4, SOX6 |
| glucose homeostasis | 4.37 | FOXA1, PDX1, HNF1A, SOX4 | —— | —— |
| positive regulation of cell cycle | —— | —— | 6.01 | FOSL1, FOXG1, TCF3, NR2E1 |
| epithelial to mesenchymal transition | —— | —— | 5.29 | TRIM28, LEF1, SNAI2, SOX9 |
| canonical Wnt signaling pathway | —— | —— | 5.26 | RARG, FOXD1, LEF1, SOX9, ISL1 |

**Fig. S2** The enriched functions and regulatory subnetworks of significantly overexpressed TFs in four cancer types. A: BRCA; B: UCEC; C: LUAD; D: LUSC; Comparison of the significantly enriched functions of TFs in BRCA and UCEC (E) and LUAD and LUSC (F). The size of TF represents the number of regulatory relations. the text box shows the GO terms significantly enriched by the TF, and the red box and text represent the common terms for the two cancers.

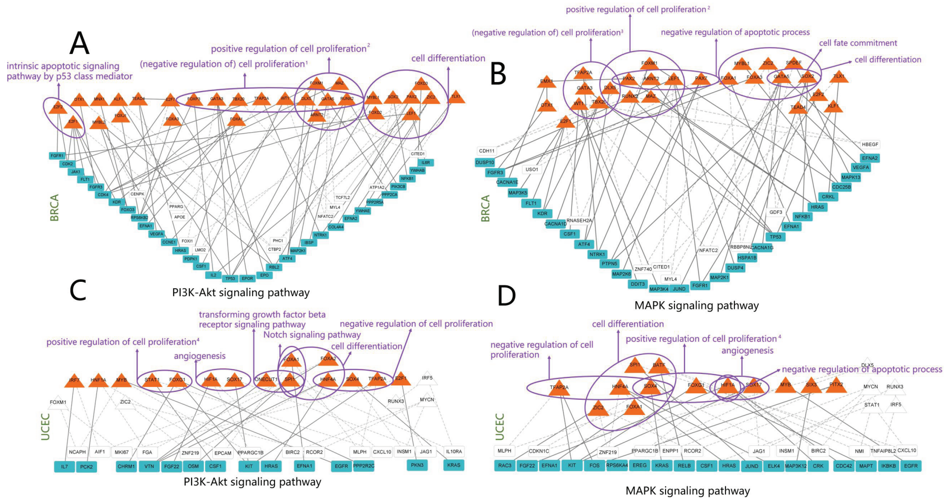


**Fig. S3** The functions and the regulatory subnetworks of overexpressed TFs regulating KEGG signaling pathways in BRCA and UCEC. A, C: The target genes belong to the PI3K-Akt signaling pathway (A: BRCA, C: UCEC); B, D: Target genes belong to MAPK signaling pathway (B: BRCA, D: UCEC).

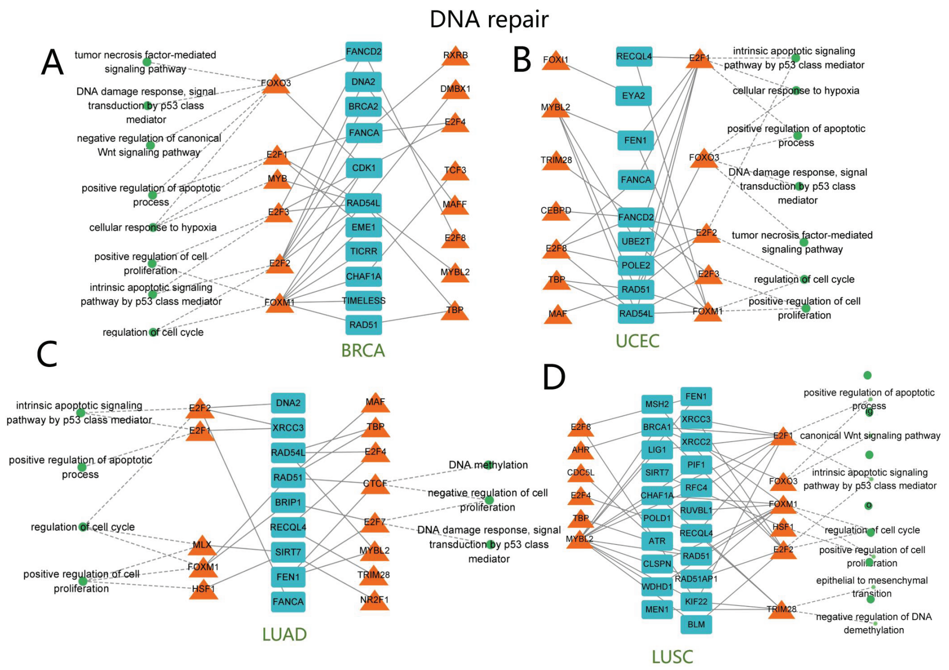


**Fig. S4** Functional subnetworks of TFs regulating overexpressed TGs in cancers. Here the TGs are involved in DNA repair. A: BRCA; B: UCEC; C: LUAD; D: LUSC. The green dots mean GO terms.

**Table S2** Statistics of enhancers related to cancer

| TCGA projects | HEDD disease types | Count |
|---|---|---|
| BLCA | urinary bladder urothelial carcinoma<br>infiltrating bladder urothelial carcinoma | 2 |
| BRCA | invasive breast carcinoma | 13 |
| COAD | colon adenocarcinoma | 0 |
| ESCA | esophageal carcinoma | 1 |
| HNSC | head and neck squamous cell carcinoma | 0 |
| KIRC | kidney cancer | 3 |
| KIRP | | 2 |
| LGG | brain glioma | 0 |
| LIHC | liver carcinoma | 155 |
| LUAD | lung adenocarcinoma | 0 |
| LUSC | lung squamous cell carcinoma | 5 |
| PRAD | prostate adenocarcinoma | 3 |
| SKCM | skin melanoma<br>cutaneous melanoma | 0 |
| STAD | stomach adenocarcinoma | 227 |
| THCA | thyroid carcinoma | 73 |

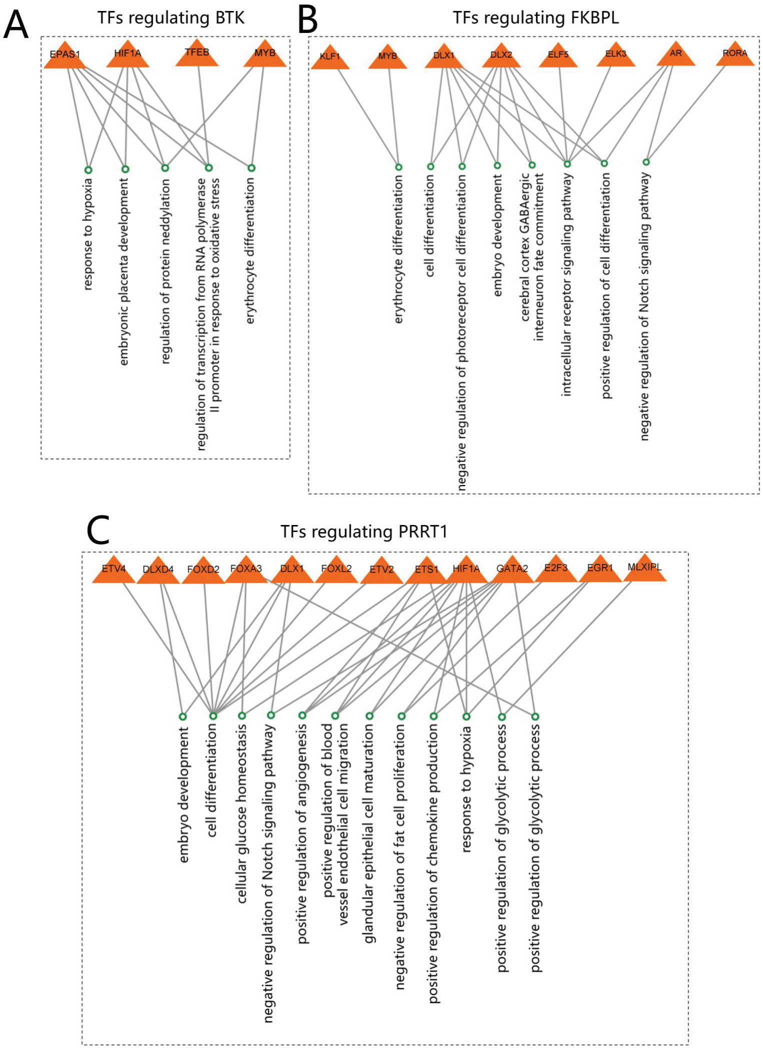


**Fig. S5** Some TFs regulating relevant oncogenes (A: FKBPL, B: BTK, and C: PRRT1) and their related biological functions.

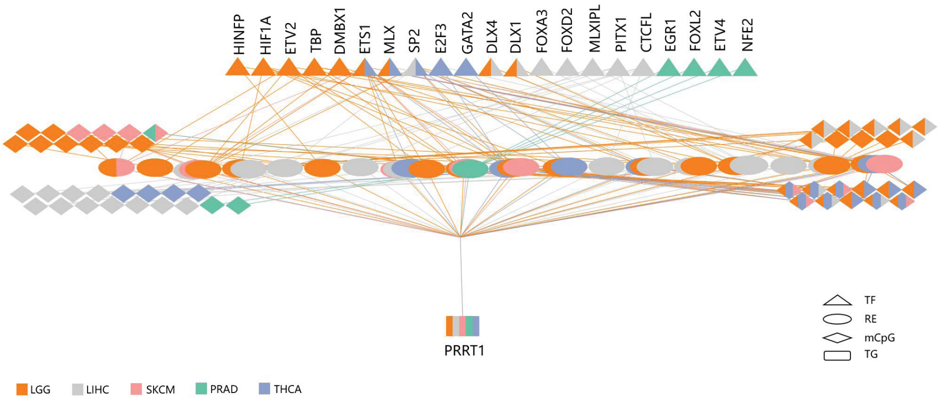


**Fig. S6** A multiplex complex regulatory relationship TF-RE(-mCpG)-*PRRT1* that regulates *PRRT1* in multiple cancers, where mCpG represents the methylated CpG site on DNA.
